# Unravelling the bottom-up and top-down control of a worldwide chestnut tree pest invader through integrative ecological genomics

**DOI:** 10.1101/2024.11.23.624965

**Authors:** Jean-Loup Zitoun, Raphaël Rousseau, Joris Bertrand, Eve Toulza, Martin Rosalie, Sébastien Gourbière

## Abstract

Biological invasions have become a major threat to all agro-ecosystems. Estimating the bottom-up and top-down forces controlling the spread of invasive insects is a key challenge to lessen their burden on crops, forestry, and biodiversity. We combined ecological, metabarcoding and population genomics analyses with an integrative modelling of the invasion of a global insect pest to identify the impacts of its chestnut tree resource, natural enemies and biological control agent in Eastern Pyrenees. The host tree frequency and genomic variation associated with common resistance pathways had effects 4-10 times greater than the native hyperparasite community on the pest’s invasion potential (R_0_). The >90% field rates of hyperparasitism by the control agent and the associated 80% reduction in pest infestation are likely to be deceptive as our modelling consistently predicts their long-term coexistence with periodic re-emergences. Such a persistent ‘co-invasion’ scenario calls for a thorough assessment of the impact of these global pest and control agent on natural forest ecosystems.

## Introduction

Biological invasions have been steadily increasing over the last two centuries (1, 2) and have become a major driver of life evolution by causing species extinction (3), homogenization of fauna and flora (4) and changes in biogeography (5). Invasive species span across a broad range of taxa (6) and are increasingly recognized as threats to the biodiversity, functioning and/or productivity of all (agro-)ecosystems (7, 8) that impair the services they provide to human societies with a staggering global economic cost of 1.288 trillion of US dollars over the last 50 years (9).

Invasive insects are known to have advert effects on crops (10), agroforestry (11), animal husbandry (12), human infrastructure (13) and health (14, 15). Their average annual cost has been estimated to reach around 10.4 billion over the last 60 years (16), which represents 40% of the worldwide economic cost of all invasive species, with three quarters of such a cost resulting from direct resource losses mostly incurred to forestry and agriculture (16). The number of non-native forest insects established outside of their natural range has indeed increased dramatically on all continents (1, 16-18) where they correspond, along with invasive pathogens, to one of the greatest threats to forestry and biodiversity conservation (14, 19). Since invasive pests are expected to keep on having profound impacts on natural and exploited forests (20, 21), to identify the determinants that impede or promote their establishment and local spread has become essential to improve risk assessment, anticipate its evolution, and quantify the efficacy of potential forest protection measures (22, 23).

The dynamic of local emergence of an invasive species typically depends on both ‘bottom-up’ and ‘top-down’ forces associated with resources limitation and natural enemies, respectively (24). Variations in plant palatability, nutritional quality, distribution and abundance are essential bottom-up factors that have long been thought to be prominent in limiting the populations of herbivores insects (25). Meanwhile, the idea that predators and parasitoids can down-regulate the herbivore populations below densities at which they have major effects on plant biomass has been stimulating plant ecology research ever since it was first proposed to provide a tentative answer to the question ‘Why is the world green?’ (26). To compare the relative importance of bottom-up and top-down forces has naturally emerged as critical to understand the population ecology of herbivores (e.g. 27-29). In a recent meta-analysis run over 350 herbivorous insect species, top-down forces were found to have about 2-fold higher impacts than bottom-up factors on abundance and life-history traits in both natural and cultivated environments (25), which provided compelling evidence of the necessity of integrative tritrophic approaches to understand the dynamic of forest invasion by non-native insects (27, 30).

The Asian chestnut gall wasp, *Dryocosmus kuriphilus*, is the most dangerous insect pest of chestnut trees (*Castanea* spp.) affecting orchards and forests worldwide (31). Native to China, this hymenopteran of the Cynipidae family was first reported to have spread to Japan in 1941 (32) and other Asian countries (33), before to be introduced in North America in 1974 (34) and in Europe in 2002, where it has invaded a wide range of regions from Italy (35). While this invasive insect pest can affect species of the genus *Castanea* and hybrid clones, there is a broad variability among clones regarding the level of infestation, suggesting a genotype-dependent variation in susceptibility (36, 37) that is thought to contribute explaining the variations in infestation observed at the individual tree level within populations of *C. sativa* (38). The individual rate of chestnut tree infestation by *D. kuriphilus* has also been shown to decrease in mixed stands, as compare to pure *C. sativa* monoculture, in line with the general ‘associational resistance’ and ‘resource concentration’ hypotheses (39, 40). Along with these typical bottom-up factors, *D. kuriphilus* recruits natural enemies that can contribute to the biotic resistance of the invaded environment. Various fungal and hymenopteran species have been shown to hyperparasite the invasive chestnut gall wasp (41), in particular parasitoids from the Cynipini communities established on oaks (42-44). However, while these native insect parasitoids are readily able to find and lay eggs into *D. kuriphilus* galls, their impact on its invasion remains limited (44), so that bio-control using a Chinese parasitoid, *Torymus sinensis*, has been intended in Japan (45), North-America (46) and Europe (47). To gain a proper quantitative understanding of the determinants of *D. kuriphilus* invasion therefore requires to measure the bottom-up and top-down forces at works, and to integrate such empirical findings into dynamical models of the tri-trophic interaction between *C. sativa, D. kuriphilus* and its native and introduced hyperparasites. While such integrative approaches have proven efficient to assess the spread of invasive plants and to anticipate the efficacy of their biological control agents (48, 49), they remain to be implemented for herbivorous insects such as *D. kuriphilus*.

In this contribution, we investigated the spread of *D. kuriphilus* in the natural chestnut tree forests of the Pyrénées-Orientales, an area located in the French Eastern Pyrenees, where the invasive pest has been informally reported since 2013, and where *T. sinensis* has been sporadically released in several places since 2014 (50). We conducted a two-years ecological field study in 23 sites located in different vegetal formations characterized by varying chestnut trees frequency to provide a comprehensive view of the local infestation of the chestnut tree populations by *D. kuriphilus*. In those sampling sites, we concomitantly implemented a normalized Genotyping-by-Sequencing approach (nGBS) of nearly a hundred chestnut trees, allowing to understand the impact of plant host genetic determinants on the level of infestation, and Genome-Wide Association Studies (GWAS) to identify loci associated with susceptibility to *D. kuriphilus* infestation. We further performed metabarcoding analyses of *D. kuriphilus* gall contents to identify their hymenopteran parasitoids, combined with emergence experiments, to estimate the rate of *D. kuriphilus* hyperparasitism by native insect and fungal species, and by the biological control agent, *T. sinensis*. Those ecological and genomic data were integrated into a dynamical model of the host-parasite-hyperparasites interactions to quantify the importance of bottom-up factors, i.e. the abundance, frequency and genetic susceptibility of chestnut trees, and top-down factors, i.e. native and introduced parasitoids, on the invasion potential of *D. kuriphilus* and its biological control by *T. sinensis*.

## Results

### Widespread and heterogeneous invasion of the *C. sativa* populations by *D. kuriphilus*

The chestnut tree populations located in the 23 sampling sites were all found infested by *D. kuriphilus* over the 2-years of our field study, which provided the first quantitative evidence of a widespread invasion of the Pyrénées-Orientales by this hymenopteran parasite that was first reported in the area in 2013 (Figs. 1-2, Table S1).

**Fig. 1.**
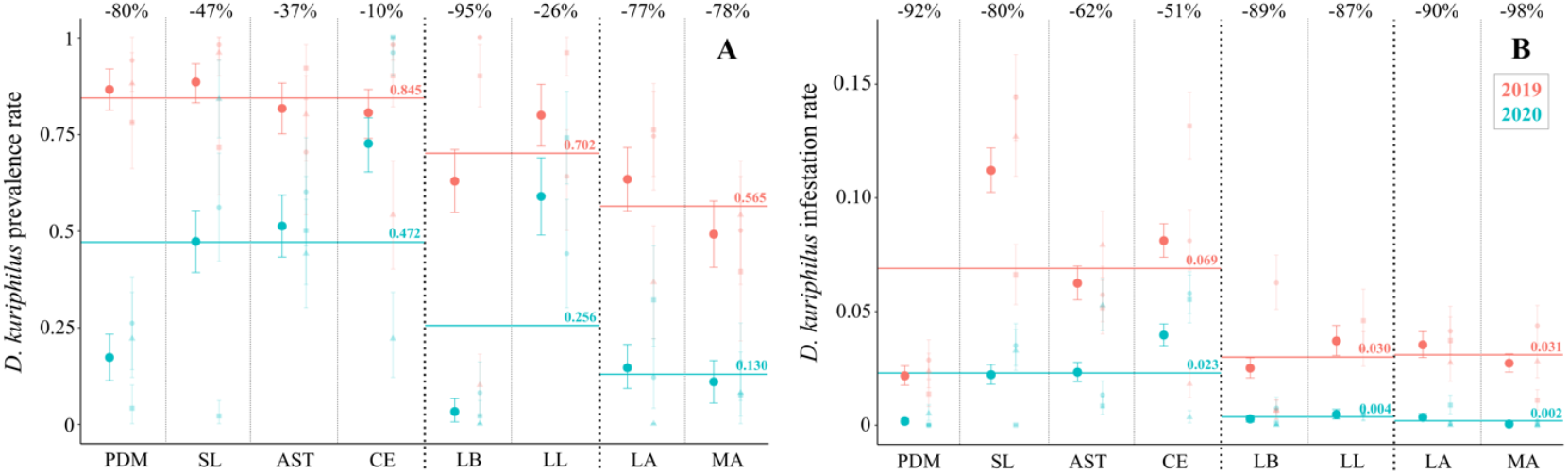
Infestation of the *C. sativa* populations by *D. kuriphilus* in the Pyrénées-Orientales. Average prevalence **(A)** and rate **(B)** of infestation and their 95% confidence intervals estimated in 2019 and 2020 appear in red and blue, respectively. Sampling sites were located in 8 stations of the Vallespir (PDM-Prats de Mollo, SL-Saint Laurent, AST-Arles sur Tech and CE-Céret), the Aspres (LB-La Bastide, LL-Llauro) and the Albères (LA-Laroque, MA-Massane) massifs (Fig. 2). For each stations, the average infestation measures are given per station and per site on the left and right hand sides of the corresponding columns. The prevalences and rates of infestation averaged across each massif are indicated by horizontal bars in figures 1A and 1B. The percentage of reduction in the average infestation measures observed between 2019 and 2020 in each station are indicated at the top of figures 1A and 1B.

In 2019, 75% of individual chestnut trees were infested with an average of 0.05 galls per leaf, representing a parasitic burden of 1 gall every 20 leaves (Table S1). The prevalence of infestation (proportion of infested trees) showed spatial heterogeneity between sites (χ^2^ = 50.6, df = 22, p = 4.9 10^−4^) and, albeit to a lower extent, between stations (χ^2^ = 13.3, df = 7, p = 6.4 10^−2^), with a larger prevalence observed in the Vallespir (0.845 ± 0.029) than in the Aspres (0.702 ± 0.06) and in the Albères (0.565 ± 0.061) (Fig. 1A). The rate of infestation (average number of galls per leaf) also showed a clear heterogeneity between sites (χ^2^ = 479.8, df = 22, p < 10^−15^) and stations (χ^2^ = 287.4, df = 7, p < 10^−15^), with similarly higher infestation in the Vallespir (0.069 ± 0.004) than in the Aspres (0.030 ± 0.004) and the Albères (0.031 ± 0.003) (Fig. 1B).

A decrease in both the prevalence and rate of infestation was observed between 2019 and 2020 across all the studied area. The overall proportion of infested trees dropped to 34% with an average of 0.013 galls per leaf, representing a reduction of 57.8% and 79.3% in those two infestation measures (Table S1). Despite significant variations between stations and sites in the amplitude of changes in infestation levels (Fig. 1A-B), the spatial patterns observed in 2020 were relatively similar to those observed in 2019. A significant heterogeneity was found at the site and station scales for both the prevalence (χ^2^ = 164.7, df = 22, p < 10^−15^ ; χ^2^ = 107.2, df = 7, p < 10^−15^) and rate (χ^2^ = 428.1, df = 22, p < 10^−15^ ; χ^2^ = 295.3, df = 7, p < 10^−15^) of infestation. The proportion of infested trees and the number of galls per leaf remained on average higher in the Vallespir (0.472 ± 0.040 and 0.023 ± 0.002) and lower in the Aspres (0.256 [0.204; 0.312] and 0.004 ± 0.001) and in the Albères (0.130 ± 0.040 and 0.002 ± 0.001). Accordingly, there was a strong correlation between the level of prevalence (ρ = 0.563, p = 4.2 10^−3^) or rate (ρ = 0.653, p=5.4 10^−4^) of *D. kuriphilus* infestation observed in the 23 sampling sites over the two consecutive years.

This field study provides clear evidences that the invasive *D. kuriphilus* has widely spread across the chestnut tree populations of the Pyrénées-Orientales with higher levels of infestation in the Vallespir than in the Aspres and the Albères and significant heterogeneities at smaller (station and site) scales. To unravel the potential ‘bottom-up’ and ‘top-down’ determinants of those spatial variations in infestation, we characterized the ecological and genetic structure of the local chestnut tree populations, and the abundance and species composition of the hyperparasite community.

### A patchy and heterogeneous *C. sativa* distribution with a low genetic structure

The distribution of chestnut trees in the Pyrénées-Orientales that we mapped using publicly available national inventories (Fig. 2A) is notably patchy and heterogeneous as they belong to different vegetal formations (IGN, 2018), with an overall density ranging from 250 to 1570 individuals per hectare, among which *C. sativa* accounted for 15.8% to 90.8%. The overall density of trees was found to decrease from the Vallespir (1055 ± 63) and the Aspres (1100 ± 65) to the Albères (753 ± 54), with higher frequencies of chestnut trees in the ‘chestnut coppice’ and ‘deciduous high forest’ of Vallespir (0.639 ± 0.08), than in the ‘high forest’ of the Aspres (0.594 ± 0.01) and in the ‘Coniferous high forest’ of the Albères (0.400 ± 0.01).

**Fig. 2.**
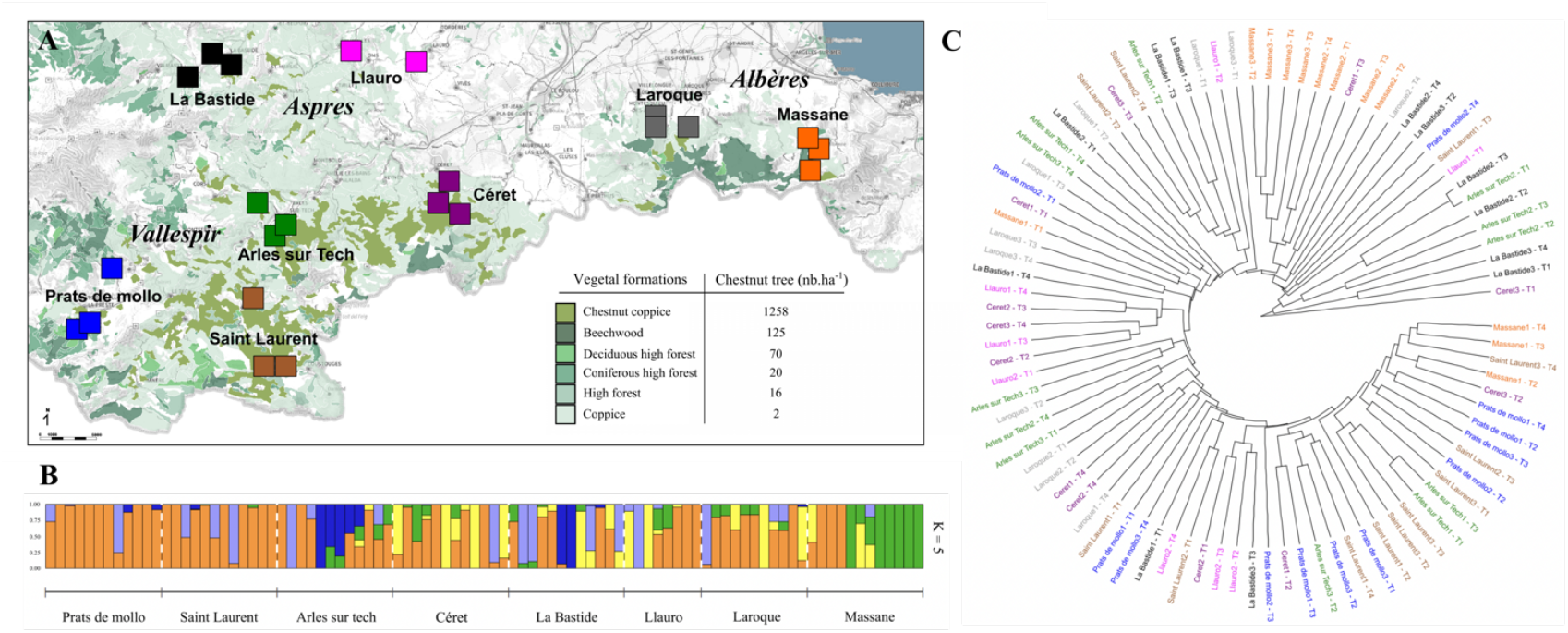
The heterogeneous distribution and low genetic structuring of *C. sativa* populations in the Pyrénées-Orientales. **(A)** Map of the different vegetal formations containing chestnut trees and the 23 sampling sites. **(B)** Genetic structure of the chestnut trees defined with respect to the 5 ancestral populations identified from the nGBS of 91 individuals, and the corresponding **(C)** phylogenetic tree, that both show a low genetic structuring of the chestnut tree populations. Colors in figures 2A and 2C are set to differentiate our 8 sampling stations, while colors in figure 2B shows the 5 putative ancestral populations identified from nGBS.

Despite such a heterogeneous distribution, the local chestnut tree populations appear to be a weakly structured genetic unit. This current structure assessed by normalized Genotyping-by-Sequencing (nGBS) using Restriction-site Associated DNA sequencing (RAD-seq) of 91 individuals located across our 23 sites was best explained as originating from 5 ancestral populations that contributed to today’s populations with no pronounced spatial pattern, although one of them was over-represented in two sites of the station ‘Massane’ (Fig. 2B) where it was first introduced for wood production at the end of Second World War. The phylogenetic tree built from the corresponding 449 markers consistently supported this lack of conspicuous spatial genetic clustering (Fig. 2C), and Jost’s *D* index confirmed the low genetic differentiation between the chestnut tree populations of the 8 stations (*D* = 0.004) and 23 sites (*D* = 0.01), despite clear genetic variations at the individual level (Fig. 2B-C).

To assess if these ecological and genetic variations contributed as ‘bottom-up’ determinants of infestation, we looked at the correlation between the prevalence and rate of infestation and the chestnut tree density, frequency and standard genetic indexes calculated at the individual and/or site level using nGBS approach.

### ‘Bottom-up’ determinants of *D. kuriphilus* infestation

A generalized linear mixed modelling (GLMM) regression analysis revealed that the frequency of chestnut tree in a sampling site strongly favors its infestation by *D. kuriphilus*. Significant positive correlations were indeed observed between chestnut tree frequency and both site prevalence and rate of infestation (Table 1A-B). Meanwhile, the density of trees in the site had no effect on the level of site infestation, reinforcing the idea that the spread of *D. kuriphilus* is modulated by the host species frequency and the local tree diversity (51) rather than by absolute densities. The site prevalence of infestation was further negatively correlated with the rate of private alleles (Table 1A), while such genetic variations had no effect on the site rate of infestation (Table 1B). On the contrary, the individual rate of private alleles was found to be associated with lower rates of infestation measured at the tree level (Table 1C). Interestingly, none of the prevalence and rate of infestation measured at the individual or site level was found to be correlated to any of the other calculated genetic indexes, i.e. nucleotide diversity, heterozygosity and inbreeding coefficient.

**Table 1.** Bottom-up determinants of *D. kuriphilus* infestation. The best GLMM models to explain the prevalence and rate of infestation at the site level (A, B) and the individual rate of infestation (C) are given with their parameters estimate and support.

| Fixed effects | Estimate | Standard error | df | P |
| --- | --- | --- | --- | --- |
| **A** Response variable : site prevalence of infestation |  |  |  |  |
| Intercept | -0.560 | 0.991 | 42 | 0.5746 |
| Chestnut tree frequency | 2.955 | 0.817 | 42 | 0.0008 |
| Site rate of private alleles | -8166.863 | 3871.641 | 42 | 0.0409 |
| **B** Response variable : site rate of infestation |  |  |  |  |
| Intercept | -4.547877 | 0.6486244 | 43 | 0 |
| Chestnut tree frequency | 1.561707 | 0.6191002 | 43 | 0.0154 |
| **C** Response variable : individual rate of infestation |  |  |  |  |
| Intercept | -3.234 | 0.593 | 181 | 0 |
| Individual rate of private alleles | -11128.909 | 4729.280 | 181 | 0.0197 |

### Variations in immune response genes correlate with *D. kuriphilus* infestation rate

A Genome Wide Association Study (GWAS) showed that Single Nucleotide Polymorphisms (SNPs) present in genes related to immune system and response to other organisms are significantly over-represented among loci identified from the nGBS method. The rates of infestation measured in 2019 and 2020 allowed to identify 33 and 491 SNPs significantly associated with those phenotypes, with 22 of them being linked with both (2019 and 2020) phenotypes and found to be distributed across 18 different loci. Those 502 SNPs were distributed in 323 different loci, 20 were identified as putative transposon/retrotransposon sequences, 75 remain uncharacterized while 228 could be identified as gene sequences, with 137 unique similar genes in *Arabidopsis thaliana* (Data S1). Gene ontology analysis revealed a significant enrichment in genes related to the response to other organisms and organic substance, (17/137), regulation of transcription (8/137), post-translational protein modification (13/137), the regulation of immune system (7/137), and to programmed cell death (7/137). Those genetic variations could explain the observed heterogeneity in individual susceptibility to *D. kuriphilus* infestation.

### ‘Top-down’ determinants of *D. kuriphilus* infestation

The dissection of 2110 galls collected across the 8 sampling stations and over the two years of our field study showed that only ∼5.9% of *D. kuriphilus* larvae developing in chestnut trees galls reached the adult stage and emerged, leaving empty lodges with typical emergence holes. The remaining lodges were found occupied by hyperparasitic insects (with 96.5% and 3.5% larvae and pupae/adults, respectively), fungi, or empty with no hole as a result of a failure of *D. kuriphilus* development (Fig. 3). The rate of insect hyperparasitism was similar in both years (∼83.5% in 2019 and ∼77.8% in 2020) and across stations (with a Fano factor of 0.003 and 0.002 in 2019 and 2020). The rate of fungi hyperparasitism was also found not to vary between years (6.7% in 2019 and 4.6% in 2020) and showed little variations between the sampled populations (with a Fano factor of 0.02 and 0.006 in 2019 and 2020). Meanwhile, the proportion of lodges where *D. kuriphilus* had failed to develop was found to be very consistent from one year to another (∼9.8% in 2019 and ∼11.7% in 2020) and between places (with a Fano factor of 0.029 and 0.013 in 2019 and 2020).

**Fig. 3.**
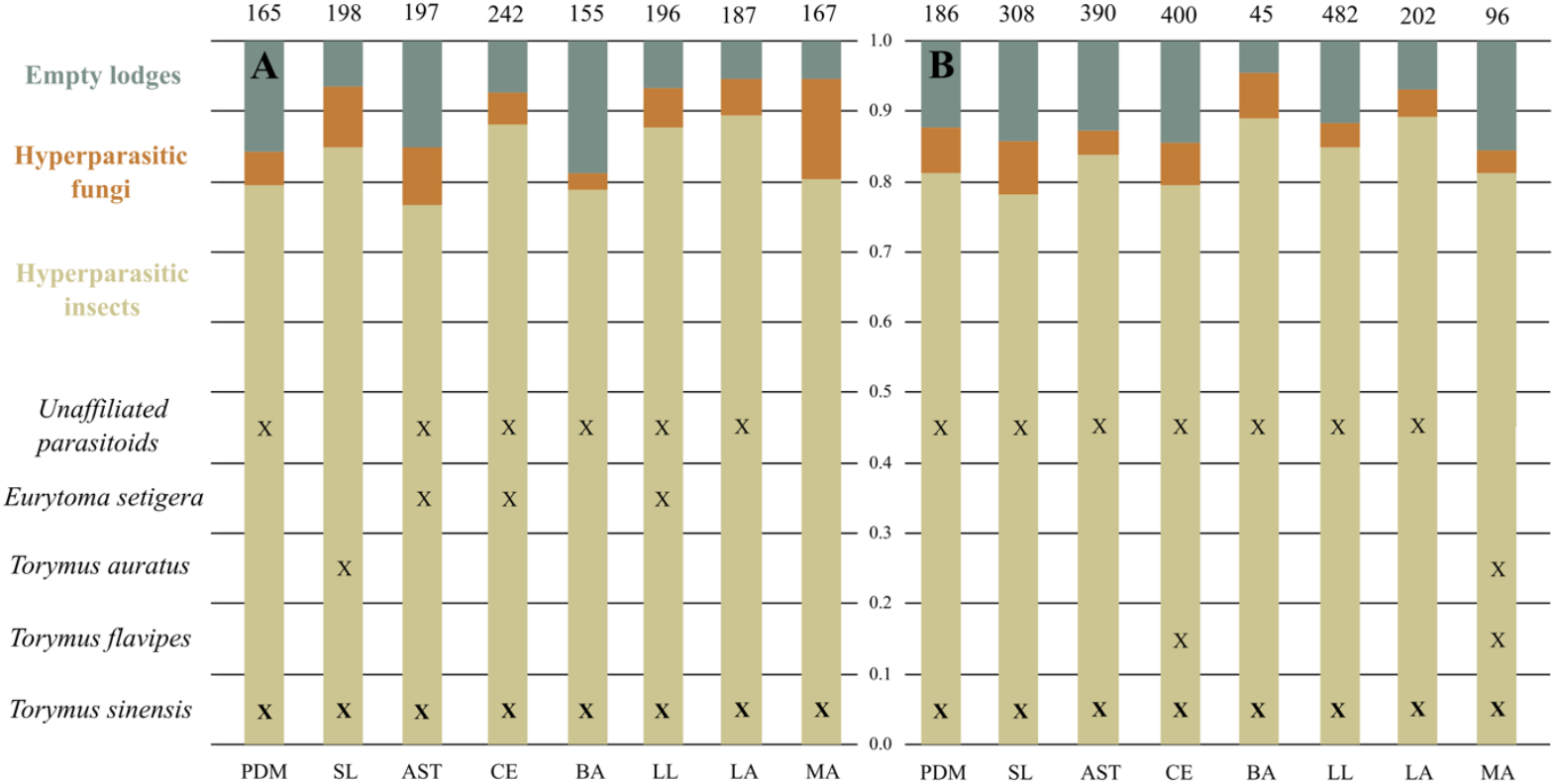
Top-down determinant of *D. kuriphilus* infestation. A total of 2110 galls were collected and dissected during our two-years field study. The corresponding numbers of lodges are indicated above each column. Among those 5.9% were found with a hole of *D. kuriphilus* emergence (not shown). The histograms indicate the proportions of the remaining lodges (with no hole of emergence) that were found with hyperparasite insects, hyperparasite fungi, and empty as a result of a failure of *D. kuriphilus* development in 2019 (**A**) and 2020 (**B**). The presence of the different identified and unaffiliated parasitoids species in each station is indicated by crosses and the bold crosses indicate the dominant species. Sampling stations are labelled as in Fig. 1: PDM-Prats de Mollo, SL-Saint Laurent, AST-Arles sur Tech and CE-Céret, LB-La Bastide, LL-Llauro, LA-Laroque, MA-Massane.

The taxonomic identification based on the ITS metabarcoding of 3002 hyperparasitic insects found in the collected galls revealed the presence of at least 6 hymenopteran species; the Asian control agent, *Torymus sinensis*, 3 native species; *Torymus auratus, Torymus flavipes, Eurytoma setigera* and at least 2 unaffiliated species (see Data S2 ans Fig. S1). Although native species varied between stations and years, the control agent was identified in all stations in both 2019 and 2020, demonstrating that it has widely spread and established itself across all the study area since its introduction in 2014. The relative abundance of *T. sinensis* among all hyperparasite species was assessed by bringing 45 galls from each of the 8 stations and following emergence in the lab. Among the 404 hyperparasites that emerged within 8 months, 386 specimens were morphologically affiliated to *T. sinensis* (95.5%) and 18 specimens were identified as native hyperparasite species (4.5%) with little variations among the 8 localities (with a Fano factor of 0.048).

Our ecological and genomic field study provided evidences that i) the frequency of chestnut trees, ii) the tree individual genetic differentiation (measured by the rate of private alleles), and iii) the presence of introduced and native hyperparasitic insects and fungi, are significant bottom-up and top-down determinants of the infestation of natural *C. sativa* populations by *D. kuriphilus*. To gain a deeper quantitative understanding of the interactions between *C. sativa, D. kuriphilus* and its hyperparasites, we tailored a population dynamic model integrating our field estimates of all the above determinants, and allowing to further explore their impacts on the regulation of *D. kuriphilus* invasion through various sensitivity analyses.

### The invasion potential (*R0*) of *D. kuriphilus* primarily depends on bottom-up determinants

Using this modelling, we identified the expression of *D. kuriphilus* invasion potential as measured by its per capita annual multiplication rate *R*_*0*_ in the natural environment (Supplementary Text S1). The *D. kuriphilus*’s *R*_*0*_ was estimated to be equal to 15.9 by using estimates of its life-history traits (Table S2), of the average frequency 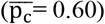 and genetic susceptibility 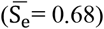 of chestnut trees, and of the average rates of infection by native hyperparasitic insects 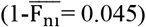 and fungi 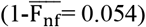 (Table S1). This invasion potential was shown to be about 1.5 more sensitive to the bottom-up than to the top-down determinants, as any 1% increase in p_c_ or S_e_ are predicted to increase *R*_*0*_ by 0.83% and 0.73%, while similar changes in 1-F_ni_ and 1-F_nf_ would reduce *R*_*0*_ by 0.52% and 0.53% (Fig. 4). The predominant effect of bottom-up determinants on *D. kuriphilus*’s *R*_*0*_ was even more evident when considering their range of observed variations between our 8 sampling stations. The changes in *R*_*0*_ induced by the observed spatial heterogeneities in pc were found to be 10.1 and 3.9 times broader than those resulting from spatial variations in 1-F_ni_ and 1-F_nf_. Meanwhile, changes in *R*_*0*_ due to spatial variations in Se were 4 and 1.5 times broader than those resulting from variations in 1-F_ni_ and 1-F_nf_. The outcomes of our sensitivity analysis of *D. kuriphilus R_0_* are thus consistent with the high spreading potential of this worldwide invasive species (42) and they provide a clear quantitative assessment of the predominance of the chestnut tree frequency and genetic susceptibility over the native hyperparasite community as main natural determinants of *D. kuriphilus* invasion.

**Fig. 4.**
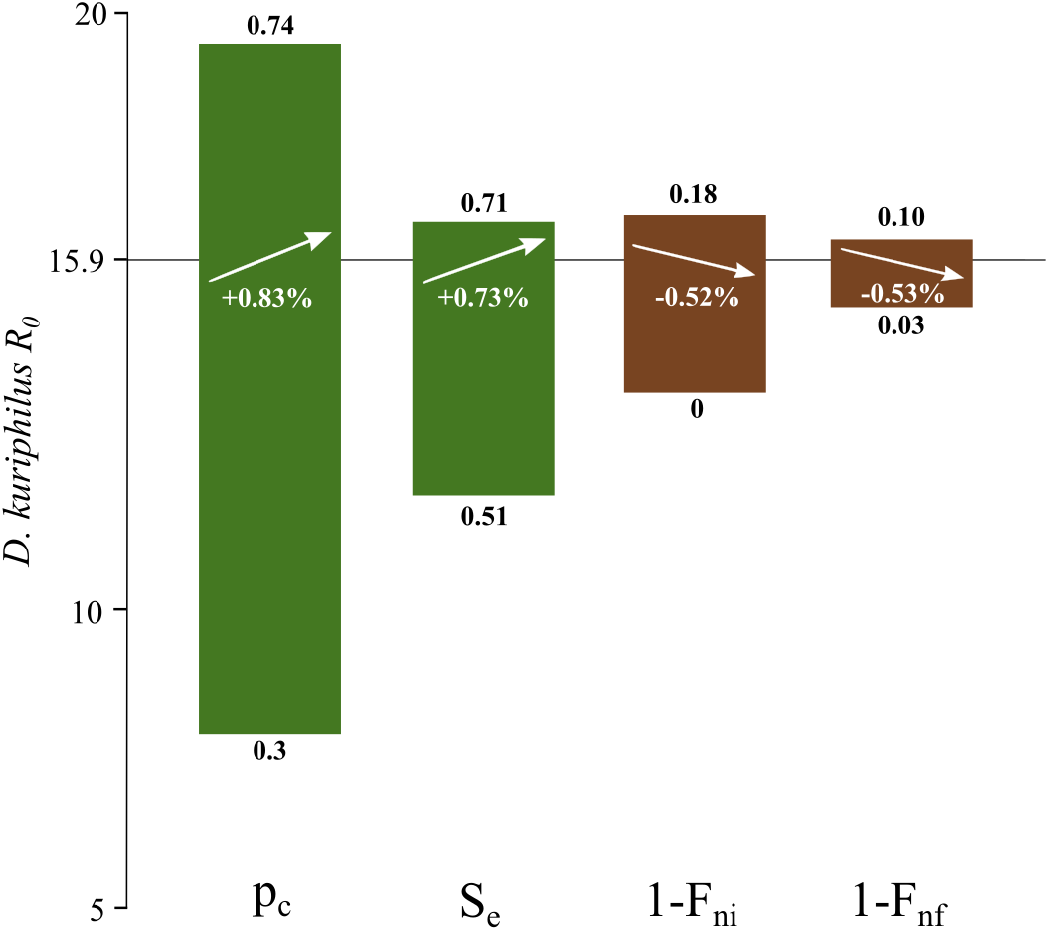
Sensitivity of *D. kuriphilus* invasion potential (*R*_*0*_) to the bottom-up and top-down determinants. The percentages of change in *R*_*0*_ in response to 1% increase in the average values of the bottom-up (p_c_, S_e_) and top-down (1-F_ni_,1-F_nf_) determinants are represented by up or down arrows. The ranges of variations in *R*_*0*_ predicted according to the maximal changes in bottom-up and top-down determinants are shown by green and brown rectangles, respectively. The maximal and minimal values of each parameter (p_c_, S_e_, 1-F_ni_ and 1-F_nf_) observed across our 8 sampling stations (Table S1) are indicated at the top and bottom of each rectangle.

### The fate of *D. kuriphilus* - *T. sinensis* interaction is set by the frequency of chestnut trees

The introduction of the commonly used biological control agent, *i*.*e*. the parasitoid *T. sinensis*, led to four different dynamics according to the frequency of chestnut trees (pc) and, to a lower extent, to their genetic susceptibility (Se). As expected, low pc or Se, typically lower than ∼0.1, does not allow for *D. kuriphilus* invasion despite the species high reproductive potential (Fig. 5A). However, as soon as pc exceeds this threshold, *D. kuriphilus* is able to invade and, whenever pc remains lower than ∼0.4, the control agent fails to get established so that the *D. kuriphilus* population growths towards its biotic capacity sets by the density of chestnut trees (Fig. 5A-B). Interestingly, in a mixed forest where pc ranges from 0.4 to 0.7, *T. sinensis* is able to spread by infesting an established *D. kuriphilus* population, which lead to dampened oscillations before a stable equilibrium is reached with both the invasive parasite and the introduced parasitoid settling down in the environment (Fig. 5A,C). Large oscillations, typically expected between a biological control agent and its target, are only predicted to occur in forests dominated by chestnut trees, i.e. when pc values are larger than 0.7 (Fig. 5A,D). Finally, none of these asymptotic dynamics were shown to depend on the size of the *D. kuriphilus* population when *T. sinensis* is introduced or on the amount of introduced individuals (Fig. 5B’-D’). This suggests that the timing of *T. sinensis* introduction during *D. kuriphilus* invasion or the release effort have no effect on the long-term outcome of the control intervention. When focusing on the range of pc and Se values observed across our 23 sampling sites, i.e. with 0.16<p_c_<0.91 and 0.427<S_e_<0.770 (Table S1), the former clearly emerged as the main bifurcation parameter that deciphers the fate of the *D. kuriphilus* - *T. sinensis* interaction dynamics between a failure of *T. sinensis* to spread, or a persistence of both species at a stable equilibrium or through large oscillations (Fig. 5A). On the contrary, the observed levels of genetic susceptibility were all large enough to have no effect on the outcome of the *D. kuriphilus* - *T. sinensis* interaction.

**Fig. 5.**
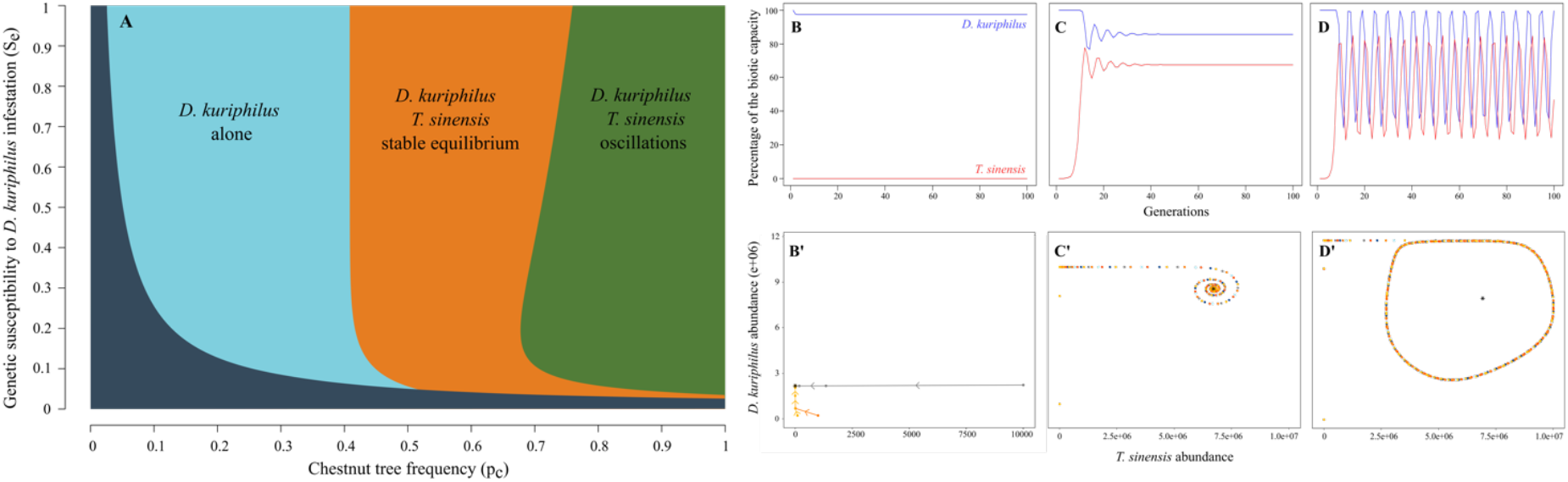
Impact of the frequency and genetic susceptibility of chestnut trees on the dynamics of interaction between *D. kuriphilus* and *T. sinensis*. **A**. Outcomes of the *D. kuriphilus* - *T. sinensis* interaction dynamics in the p_c_-S_e_ parameter plane. Dark blue = both species fail to spread, light blue = *T. sinensis* fails to spread and to control *D. kuriphilus*, orange and dark green = stable equilibrium and stable oscillations between *D. kuriphilus* and *T. sinensis*. **B-D**. Chronics describing the temporal dynamics of *D. kuriphilus* (H(t), blue line) and *T. sinensis* (P(t), red line) observed in light blue, orange and green parts of A. Both abundances are given as percentage of the species biotic limit estimated for a 1 km^2^ area. **B’-D’**. Phase planes illustrating the lack of sensitivity of the asymptotic dynamics to the initial conditions H(0) and P(0), i.e. the abundance of *D. kuriphilus* and *T. sinensis* when the latter is introduced. Each colour corresponds to a different pair of H(0) and P(0) values.

### Biological control is optimal in chestnut tree dominated forests with low susceptibility

We explored further the efficacy of *T. sinensis* release in forest environments characterized by different frequencies and genetic susceptibilities of chestnut trees by predicting the percentage of the *D. kuriphilus* population that could be removed by the introduction of *T. sinensis* (Fig. 6). In line with the dynamical outcomes shown in Fig. 5A, forests with too low frequency of chestnut trees and/or chestnut trees with too low genetic susceptibility do not allow for the spread of the control agent because of the too scarce abundance of *D. kuriphilus*. Once the forest environment allows for a stable equilibrium between *D. kuriphilus* and *T. sinensis*, the rate of control of the invasive by the biological agent rapidly increases with the genetic susceptibility of chestnut trees until it reaches an optimal value that is typically larger than 50% when the frequency of chestnut trees is above ∼0.7. Meanwhile, the efficiency of biological control slowly decreases as soon as the genetic susceptibility of chestnut trees exceeds 0.05-0.15, although it remains larger than 25% for any forests with a frequency of chestnut trees over ∼0.8. Accordingly, the biological control agent is expected to be more efficient in chestnut tree dominated forests exhibiting large oscillations in *D. kuriphilus* and *T. sinensis* abundances, than in mixed-forest where the interaction dynamics converges towards a stable equilibrium.

In the forest environment typically encountered in the Pyrénées-Orientales, i.e. with 0.16<p_c_<0.91 and 0.427<S_e_<0.770 (Table S1), *T. sinensis* is predicted to lower the abundance of *D. kuriphilus* by up to 44%. In such conditions, the efficiency of biological control is expected to be primarily determined by the proportion of chestnut trees stands since genetic susceptibility no longer is a limiting factor in the flattening part of the relationships shown in Fig. 6.

**Fig. 6.**
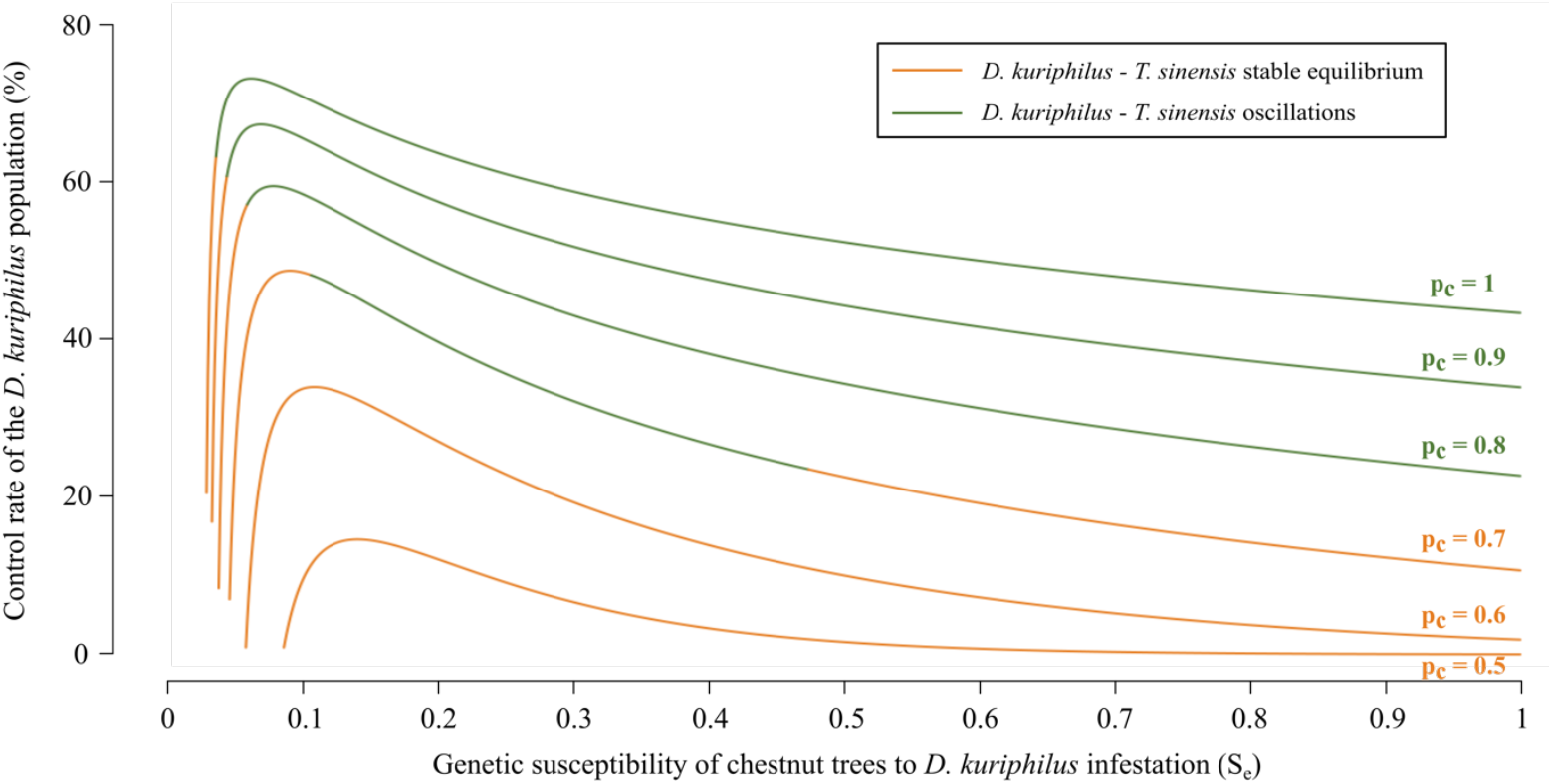
Efficacy of *D. kuriphilus* biological control in different forest environments. The efficacy of control is measured as a percentage of reduction in *D. kuriphilus* abundance following the introduction of *T. sinensis*. The abundance of *D. kuriphilus* with and without *T. sinensis* was evaluated numerically after a stable (orange) or oscillatory (green) equilibrium was reached, as shown in Fig. 5.

## Discussion

This ecological field study provided the first evidence that the chestnut tree pest, *D. kuriphilus*, has widely spread in the Pyrénées-Orientales area, thereby contributing to the global effort of documenting its worldwide invasion. The levels of infestation recorded in the first year were relatively high, with 75% of infected chestnut trees and 30% of our sampling sites exhibiting infestation rates that exceeded the threshold of 0.6 galls per bud above which *D. kuriphilus* can induce drastic decrease of tree productivity (36). Such high infestation levels reported 6 years after the first vernacular report of this invasive species in the area are consistent with those observed after a similar amount of time following its arrival in Switzerland and Italy (52). Decreases of ∼60% and ∼80% of the prevalence and rate of infestation were observed in 2020, with no site remaining highly infested and 80% of sites falling below a threshold of 0.3 galls per bud, where they are considered as weakly infested with no substantial threat for tree production or growth (36). Behind this clear temporal pattern, heterogeneities in infestation were observed at site and individual tree levels, which we exploited to uncover the bottom-up and top-down determinants of *D. kuriphilus* invasion.

We first aimed at characterizing the bottom-up factors that could shape the spatial variations observed in *D. kuriphilus* infestation. Data from the national forest inventory (53) showed that *C. sativa* is one of the most abundant species in the area with a patchy distribution spread in different plant formations where its frequency varies between 16% and 91%, which provided a truly heterogeneous landscape for *D. kuriphilus* invasion. This fragmented distribution is associated with a low genetic structuring between localities separated one from another by distances ranging from 7 to 46 km. Still, the overall chestnut tree population appears to be constituted of 4 indigenous strains and an additional exotic variety introduced in ‘La Massane’ for exploitation and reforestation purposes, with noticeable events of hybridization between strains. Interestingly, our analysis of the impact of those spatial variations in the plant resource showed that a high chestnut tree frequency and a low rate of private alleles are associated with higher levels of prevalence and rates of infestation at the site level. The identification of resource availability as a bottom-up factor facilitating the invasion of *D. kuriphilus* in forest environments is consistent with previous evidences that infestation rates (40) and pest-induced levels of defoliation (51) are higher in chestnut tree monocultures than in mixed stands, and with a broad range of studies highlighting the primary importance of the access to the trophic resources for the spread of invasive herbivores (17) and plant parasites (54). While significant variations in the number of *D. kuriphilus* eggs and galls per bud were previously observed among European and Asian *Castanea* species and their hybrids (37, 55) and between local ecotypes in Southern Italy (56), our results provide the first evidences that *C. sativa* genetic variations can also significantly affect the spread of *D. kuriphilus* across the natural forest environment. Such findings about the roles played by both chestnut tree abundance and genetic diversity in the spread of their worldwide invasive parasite through the forest environment concur with the ecological theories predicting that resources in high frequency shall be most vulnerable to parasites and pathogens (‘resource concentration’ hypothesis, (39, 57)) and that their genetic homogeneity is favorable to diseases spread (‘disease-diversity’ relationship, (58)). Meanwhile, our ecological field study further led to observe that, even in highly infested sites, some chestnut trees remained uninfected or infected at strikingly low levels, which was ultimately corroborated by analyses performed at the individual level that showed a negative correlation between infestation level and the rate of private alleles. According to Jiang et al. (59), such individual genetic variations are primarily expected to arise among trees that are not part of the indigenous population or that have experienced outcrossing with introduced strains. Our results thus suggest that the introgressions between the coexisting endemic and exotic strains observed from the nGBS approach, are likely to have provided local chestnut trees with greater resistance to *D. kuriphilus* invasion, as typically expected from the novel agricultural practice of intercropping proposed to counter the effects of monoculture (60). A genome wide association study then allowed to identify genes likely involved in the chestnut tree resistance to *D. kuriphilus* as they are associated with biological processes and protein families present in the hypersensitive response pathway that mediates host responses upon pathogen infection. Significant SNPs were over-represented in genes that can act in elicitor recognition, activation of the transcription, salicylic and jasmonic acid biosynthesis, transport and regulation, and programmed cell death through hypersensitive response. The identification of these genes is consistent with the previous report of an hypersensitive response in the totally resistant chestnut tree strain ‘Bouche de Bétizac’ (61) and the observation of a differential expression of leucine rich repeat proteins, receptor-like protein and kinases, cellular death related genes, transcription factor and miRNA target genes when comparing susceptible and resistant chestnut tree varieties that were infected or not by *D. kuriphilus* (62, 63). Our GWAS analysis therefore allowed for the identification and partial reconstruction of the genetic response pathway activated by chestnut trees while facing invasion by their worldwide insect pest. Such a cost-effective approach, based on Restriction site Associated DNA Sequencing of less than a 100 trees, offers a promising way to unravel the key genomic determinants shaping the interactions between non-native insect pests and tree species even in the absence of host reference genomes as already suggested by Pais et al. (64).

While characterizing the bottom-up factors influencing spatial heterogeneities in *D. kuriphilus* infestation, we concomitantly investigated the role of the top-down control exerted on the invasive pest by native and introduced parasitoids. Native hyperparasite species were collected in galls formed by *D. kuriphilus* on chestnut trees, and their molecular taxonomy allowed to identify at least 5 hymenopteran species infesting *D. kuriphilus* larvae in the Pyrénées-Orientales (Data S2 ans Fig. S1). The identified species were generalist parasitoids known to preferentially infest oak gall-forming parasites (42) that were previously observed in Cynips galls formed on chestnut trees in other parts of France (50). Although such native parasitoids were obviously able to adapt their diet to the new resource opportunity, their rate of parasitism of *D. kuriphilus* larvae remained low (4.46%), which is also consistent with previous field estimates obtained in neighbouring areas in south of France (50) and elsewhere in Europe (65, 66). We observed a concomitant hyperparasitism of *D. kuriphilus* by fungal species at a rate that was similarly low (5.44%) and in agreement with previous reports of an average rate of *D. kuriphilus* larvae by *Fusarium proliferatum* of 3.55% in Italy (67) and 3% to 6.8% in the US (41). The low observed rates of *D. kuriphilus* hyperparasitism by both hymenopteran and fungal endemic species clearly suggest, in line with the ‘Enemy Release Hypothesis’ (68) and with previous statements (42, 44), that the spread of this worldwide chestnut tree pest is loosely affected by natural enemies. The metabarcoding and morphological identification of the insects found in the *D. kuriphilus* galls collected in our 24 sampling sites provided strong evidence that the biological control agent, *T. sinensis*, has widely spread across all the Pyrénées-Orientales since it was first release in 2014 (50). The parasitoid was detected during both years of the study and in all 24 sites, with rates of *D. kuriphilus* parasitism that reached 92.8% in 2020. These important levels of infestation, 6 years after the introduction of the control agent, are highly consistent with the >90% rates of infestation by *T. sinensis* that were observed 5-7 years after the implementation of biological control in Italy (69), and are likely to explain the strong reduction in *D. kuriphilus* abundance observed between 2019 and 2020. Although the presence of *T. beneficus* was suggested by our metabarcoding analysis (Data S2), this species cannot accurately be discriminated from *T. sinensis* since they share the same ITS2 genotype (70), as further evidenced by the phylogenetic tree of OTUs sequences (Fig. S1). Thus, since there is no introduction record of this Japanese species in the Eastern Pyrenees, all OTUs sequences are most likely to belong to the introduced *T. sinensis*.

Although our ecological and genomic analyses unraveled the bottom-up and top-down factors influencing *D. kuriphilus* infestation levels in the Pyrénées-Orientales, we designed eco-genomic models to integrate further their corresponding estimates and provide quantitative insights into the dynamical interplay between those factors during the insect pest invasion. Our modelling allowed to derive an original expression of the invasion rate of *D. kuriphilus* (*R0*) accounting for all naturally occurring factors, and the resulting estimate of this yearly population growth rate was found to be high, i.e. 15.9, as expected given the species strong reproductive potential associated with parthenogenesis and the favourable conditions encountered, i.e. large average frequency (60%) and genetic susceptibility (68%) of chestnut trees and a low rate of infection by native hyperparasitic insects (4.5%). Importantly, the sensitivity analysis to local variations around those average field estimates showed that bottom-up determinants, and especially the frequency of chestnut trees in the forest environment, had 1.5 to 10.1 times more impacts that native parasitoids on this pest invasion rate. This first quantitative assessment of the relative impact of bottom-up and top-down factors on *D. kuriphilus* invasion sharply contrasts with the conclusion of a recent meta-analysis that top-down factors had 1.3 more effect than bottom-up factors on specialist gallers (25). As this meta-analysis primarily dealt with studies of insect herbivores within their native range, such a discrepancy provides another neat evidence of the importance of escaping natural enemies in invasion success (71) and is consistent with the large body of field observations that *D. kuriphilus* has indeed escaped most of its enemies while spreading across Asia (72), into Europe (44, 50, 65, 66) and the US (41). Our modelling made another key prediction that, following invasion and the introduction of the control agent *T. sinensis*, their interaction should, in conditions encountered in the Pyrénées-Orientales, lead to a long-term coexistence. Noteworthy, whether the two species persist around an equilibrium point or through stable fluctuations was shown to primarily depend on the frequency of chestnut trees, with a bifurcation value of ∼70% whatever be the genetic susceptibility of the host resource. Such theoretical findings encompass the prediction of host-parasitoid cycles made for a pure chestnut tree stand by the unique available model of *D. kuriphilus* - *T. sinensis* interaction (73, 74) and expanded it into a complete range of chestnut tree frequency and genetic susceptibility. As already suggested in Paparella et al. (73), the prediction of persistent oscillations echo the observation made in Japan of fluctuations between *D. kuriphilus* and its control agent, *T. sinensis*, leading to three successive resurgences of the parasite population over a 25 years period. Meanwhile, our modelling further showed that when integrating local eco-genomic estimates of naturally occurring bottom-up and top-down determinants of *D. kuriphilus* demography, its coexistence with the control agent becomes feasible in conditions typically met in Europe, which contrasts with previous conclusions (73). The main cause for such an apparent discrepancy is that, accounting for our field estimates of the effects of the genetic susceptibility of chestnut trees and of developmental failures on *D. kuriphilus* eggs survival, the rate of pest overwintering was reduced by 64% in our study, which, in line with the bifurcation analysis provided by Paparella et al. (73), lowers the amplitude of the oscillations and allows for the long term persistence of the invasive and its control agent. Our integrative modelling therefore bring a comprehensive quantitative understanding of the *D. kuriphilus* - *T. sinensis* interaction in Europe as it predicts both a strong reduction of *D. kuriphilus* in the first few years after the introduction of *T. sinensis*, as empirically shown in this study and many European countries (see (42) for a review), and the long-term persistence of these two exotic species, which complies with the field observations that *T. sinensis* has become established in the natural environment in Spain (75), France (76) and Italy (77), from which it has spread into Slovenia, Croatia, Hungary (78), Bosnia and Herzegovina (79).

We lastly used our modelling to evaluate the efficacy of biological control in such a context of long-term persistence between *T. sinensis* and its targeted insect pest. The control agent was predicted to be most efficient in reducing the abundance of *D. kuriphilus* in forest with over 70% of chestnut trees, where the interaction between the two species lead to fluctuating dynamics. The overall reduction rate of the pest population then reached 30% to 60% for typical level of genetic susceptibility estimated to range from 0.4 to 0.8, and increased with the amplitude of the oscillations as those allows for larger drop in *D. kuriphilus* abundance that can then be reduced by a factor of up to 10^−3^. Although such levels of biological control are not expected to lead to the extinction of the insect pest, they can actually drive the rate of infestation below the threshold of 0.3 galls per buds that is associated with no substantial threat for tree production or growth (36). Interestingly, in mixed forest with frequency of chestnut trees lower than 70%, the interaction is more stable and the reduction of *D. kuriphilus* abundance by *T. sinensis* remained typically lower than 20%. These predictions confirm previous conclusions that the biological control of this insect pest typically thought for chestnut trees monocultures, is likely to be much less efficient in large heterogeneous forest environments (74), albeit for a different reason. While the lower control efficacy to *T. sinensis* was there attributed to the higher dispersal abilities of *D. kuriphilus* allowing to escape in a spatially structured environment, our study shows that the change in bottom-up factors between orchards and mixed-forest patches can readily explain the outcome of the *D. kuriphilus* - *T. sinensis* interaction at a local scale.

To conclude, this integrative eco-genomic modelling study illustrates the genuine interest of implementing tri-trophic approaches to understand the dynamic of forest invasion by non-native insects (25, 30). Our combined ecological, genomic and dynamical modelling analyses provided clear evidences that the emergence of the worldwide invasive *D. kuriphilus* in the chestnut tree forests of the Pyrénées-Orientales has been predominantly shaped by bottom-up factors until the introduction of *T. sinensis*. They further demonstrated that the strong initial reduction of the pest abundance due to the control agent is unlikely to lead to the co-extinction of the two exotic species, but instead that their interaction in the Pyrénées-Orientales is entering a similar long term dynamic as those observed in Japan with periodical re-emergence of the parasite (73). Given the rising evidences that *T. sinensis* has been able to spread through natural forests in Europe (75, 78, 79), our findings clearly call for deeper assessments of the ability of the control agent to parasite native cynipidae (80, 81) and to compete (82, 83) and hybridize (84, 85) with closely related native parasitoid species.

## Material and Methods

### Dryocosmus kuriphilus life-history and its bottom-up and top-down determinants

The worldwide invasive, *D. kuriphilus*, is an Asian wasp inducing the formation of galls on stem and leaves of chestnut trees, which causes up to 80% chestnuts production loss (86) and makes infested trees more vulnerable to other parasites, most notably the pathogenic fungus *Cryphonectria parasitica* responsible for the chestnut blight (87). This hymenoptera species is univoltine, semelparous and parthenogenetic. Females emerging from galls in June-July are 1-7 days short-lived and fly to lay their asexually produced eggs in chestnut tree buds during a 2 to 3-weeks period, before to die. Oviposition can trigger a local tree hypersensitive immune response, with the accumulation of reactive oxygen species, resulting in programmed cell death and leading to the death of deposited eggs (61). The efficacy of this response shows substantial variations between chestnut tree varieties, with levels of susceptibility to bud infection varying by one to two orders of magnitude (36). Eggs escaping such a response and intrinsic developmental failures hatch within a month to develop into first instar larvae that enter an overwintering dormant stage. In spring, larvae develop into subsequent larval stages and induce the formation of galls where they eventually become pupae and adults emerging to lay the next generation of eggs. Meanwhile, hymenopteran parasitoid species target the newly formed galls to lay their progeny that will ultimately feed on *D. kuriphilus* larvae, thereby downregulating the invasive population. Such hyperparasites can include native species, although their reported parasitism rates are usually low (66, 83), together with the introduced Asian control agent, *Torymus sinensis*, an univoltine and semelparous Hymenoptera of the *Torymidae* family (47). Galls have been repeatedly shown to be further infected by local fungus that induce an additional mortality of *D. kuriphilus* larvae (41, 67). All the above bottom-up and top-down factors influencing *D. kuriphilus* life-history ultimately determine its ability to spread and succeed in a local invasion.

### Distribution of the C. sativa populations and sampling sites in the Pyrénées-Orientales

We investigated the spread of *D. kuriphilus* in the *C. sativa* populations of the Pyrénées-Orientales, a French department located on France’s Mediterranean coast along the Spanish border, in 2019 and 2020 (Fig. 2A). This part of Eastern Pyrenees is made up of highly contrasted landscapes spanning from the Vermeille Coast to the 2784 meters high Canigó Massif. The spatial distribution of the local *C. sativa* populations shown in Fig. 2A was characterized by combining i) cover maps of the different vegetal formations that were *derived* from aerial *photointerpretation* of the study area (88), with ii) estimates of the density of chestnut trees in each of these vegetal formations that were obtained through systematic clustered sampling during national forest inventories (53). The chestnut tree populations are mostly found in the Vallespir, Aspres and Albères massifs, where we based our 24 sampling sites in 8 stations; Prats de Mollo, Saint Laurent, Arles sur Tech and Céret (Vallespir), La Bastide and Llauro (Aspres) and Laroque and La Massane (Albères).

### Infestation of C. sativa populations by D. kuriphilus in the Pyrénées-Orientales

The invasion of the local chestnut tree populations by *D. kuriphilus* was assessed by two typical measures of parasitism in each sampling site. The prevalence of infestation, defined as the proportion of chestnut trees infested by at least one gall, was estimated on 50 trees in each site. The rate of infestation, defined as the mean number of galls per leaf, was estimated on 5 geolocated chestnut trees per sampling site with a sampling effort of 250 to 500 leaves per tree. The existence of heterogeneity between sites or between stations in the prevalence or rate of infestation was tested by Pearson’s chi-squared tests with Yates’s correction for continuity. Exact confidence intervals for both measures of infestation were calculated from the binomial distribution set according to the corresponding sampling effort. The correlation between the prevalence or rate of infestation measured over the two consecutive years of our field study was assessed using Kendall’s rank correlation test.

### Characterization of the C. sativa structure in sampling sites

The chestnut trees distribution in the forest stands was characterized by sampling 1000m^2^ in each of our sampling sites and by estimating their abundance and frequency among the tree community. Confidence intervals for the abundances and frequencies were drawn from Poisson and Binomial distributions, respectively.

### Chestnut trees genome sequencing and SNP calling

We randomly selected 4 out of the 5 geolocated chestnut trees in each site, from each of which a 3 cm^2^ leave sample was collected and sent in 70° alcohol to LGC Genomics GmbH (Berlin, Germany) for single-end Restriction-site Associate Sequencing (RAD-seq) using the Apek1 restriction enzyme, and following a normalised Genotyping-by-Sequencing approach (nGBS). Libraries were generated on an Illumina NextSeq 500 with an average number of ∼3 millions reads (75bp) per sample (Table S3) that were deposited in the NCBI Sequence Read Archive (SRA) under the BioProject accession number PRJNA1111809. High-quality clean reads were mapped onto the closest phylogenetically related *Castanea* reference genome, i.e. the *Castanea mollissima* genome (GCA_000763605.2) using BWA-MEM (0.7.16a-r1181) with default settings. The `ref_map.pl` module of the STACKS program (89) was used to call SNPs in each sample, based on 635818 loci with a mean coverage of 30X according to the alignment positions provided for each read.

### Genetic structure of the C. sativa populations

The genetic structure of the chestnut tree populations was established from SNPs shared by 100% of the individuals and obtained from the `populations` module of the STACKS program with default parameters, as such a stringent filter was thought to provide optimal estimates of phylogenetic distances between individuals. A poorly sequenced individual (from the second site located in Laroque, i.e. Laroque 2 - T3, see Table S3) was removed at this stage, which increased by a factor of 10 the number of SNPs shared by all remaining individuals and rose the power of our genetic structure analyses. Subsequent analyses made on the 449 resulting SNPs were achieved using R.4.1.1. and dedicated packages (90), but for the calculation of the Jost’s differentiation index (Jost’s *D*; (91)) that was done using Genodive (92). The individual admixture coefficients were inferred using the sNMF function of the LEA package (93) with K (number of ancestral populations) varying from 1 to 10, with 10 replicates for each. The SNPRelate package (94) was then used to build phylogenetic trees from the individual genomes. The initial analyses ran on the (4×24)-1= 95 chestnut trees showed that individuals from the third sampling site located in Llauro were genetically identical and very divergent from the rest of the individuals (Fig. S2), so that they were treated as outliers and the site was removed from all analyses appearing in Figs. 1-2 and Table 1. Accordingly, the final analyses of the genetic structure of the chestnut tree populations appearing in Fig. 2 were run on the remaining (4×23)-1 = 91 individuals, while genetic indexes were estimated on the 92 individuals of the 23 sites kept in the analysis (see below).

### Genetic indexes of the chestnut tree populations

We estimated standard genetic indexes using 31736 SNPs shared by at least 80% of the 92 individual genomes sampled from the 23 sites kept in the analysis. Nucleotide diversity (π), heterozygosity (*H*), inbreeding coefficient (*F*) were all estimated using the `populations` module of the STACKS program (89), but the inbreeding coefficient at the individual level was calculated with VCFtools (95). The number of private alleles was estimated for each individual and site from the total number of SNPs using the `populations` module of the STACKS program with default parameters. The frequency of private alleles was then calculated as the ratio of the number of private alleles to the total number of bases sequenced per individual or site.

### General linear mixed modelling of the bottom-up control of D. kuriphilus infestation

To identify the potential determinants of a bottom-up control of *D. kuriphilus* invasion, we searched if the observed levels of infestation could be explained by the set of ecological (overall density of trees, frequency of chestnut trees) and genetic (nucleotide diversity, heterozygosity, inbreeding coefficient and frequency of private alleles) variables that were measured on the 23 sampling sites and 92 individual trees. Statistical analyses were performed through 3 independent generalized linear mixed modelling (GLMM) to explain variations in (i) the prevalence of infestation between sites, (ii) the rate of infestation between sites, and (iii) the rate of infestation between individual trees (iii). Each statistical model best explaining those infestation levels was identified by following the procedure recommended by Zuur et al. (96). First, we characterized the impact of the sampling year on the infestation measures through a standard GLM analysis (Infestation: Estimate = -1.421, p = 3.65 10^−4^, Prevalence: Estimate = -1.7292, p = 7.99 10^−5^), in order to account for such a temporal structure (by incorporating sampling year as a random factor) in our subsequent GLMM analyses ((96), p. 323-332). Second, we identified collinearity between ecological and genetic explanatory variables using the corvif function of the AED package, and subsequently excluded the most correlated variables with a cut-off of 3 ((96), p. 387). Once heterozygosity had been removed in such a way, all our explanatory variables appeared independent one from another, with all collinearity measures ranging between 1 and 2.5. Third, the best statistical model was identified by sequentially removing non-significant predictors according to the outcomes of student tests comparing the model fitting with and without each of the remaining explanatory variables ((96), p. 220-222). All 3 GLMM analyses were conducted with R 4.3 using the glmmPQL function from the MASS package and a quasi-binomial distribution, after the original rate of infestation (i.e. the number of galls per leaf) was converted into a proportion of leaves with one gall since we never counted more than one gall on any of the >45000 observed leaves.

### Genome wide association studies

Two genome wide association studies (GWAS) were performed on the 92 genotyped chestnut trees in order to identify the genetic bases involved in the susceptibility of *C. sativa* to *D. kuriphilus* infestation. The association between the 31736 SNPs shared by 80% of the individual chestnut trees, and each of the 2 individual phenotypic measures of infestation, i.e. the rate of infestation measured on individual trees in 2019 and 2020, were calculated using the Latent Factor Mixed Models (LFMM) package in R 4.3. The significant SNPs were selected by applying a cut-off equals to ∼3.1e-05 (1/31736). The genomic regions containing these SNPs were then identified by a sequence homology analysis using *C. mollissima* as reference genome. When SNPs were located in genes described in the annotation file of the *C. mollissima* genome, we extracted the entire gene sequence. A BLASTX analysis was then performed for the sequence of each of these genes using the Plaza Dicots 4.5 database, allowing for genes to be identified when the p-value of the alignment was found ≤1e-40. When the alignment did not meet such a criterion, we used the less specific NR NCBI database and genes were identified when the alignment score was ≥200, corresponding to a p-value of 1e-45. When multiple alignments with a score ≥ 200 were found, we selected the alignment with the closest location to the SNPs being considered. For the remaining SNPs that were not found in genes described in the *C. mollissima* genome annotation file, we extracted 3000bp of the *C. mollissima* genome on each side of the SNPs, resulting in a 6001bp sequence. Sequence homology analysis was then performed using the NCBI database as describe above. Once all genes were identified, their functions and *Arabidopsis thaliana* homologs were determined using the Uniprot database (97). Finally, a singular enrichment analysis was performed using agriGO (98) with the *A. thaliana* TAIR9 database to identify a potential enrichment in some biological process among the candidate genes.

### Collection and barcoding of D. kuriphilus and its hyperparasite community

Only a fraction of the *D. kuriphilus* eggs laid in *C. sativa* buds emerge as adults because of developmental failures and hyperparasitism. The *D. kuriphilus* larvae can indeed be parasited by native hyperparasitic insects or fungi, and by *Torymus sinensis*, the hymenoptera species used as a biological control agent. To assess the importance of these potential top-down determinants of *D. kuriphilus* infestation, we combined three complementary experimental approaches. First, we dissected 770 and 1340 galls collected from each of our 8 sampling stations in 2019 and 2020, respectively. For each gall, we counted the number of lodges that were (i) empty with a hole (indicative) of emergence, (ii) empty without such a hole, or occupied by an hyperparasitic (iii) (native or introduced) insects or (iv) fungi. These allowed to estimate in each sampling station; the proportions of larvae that (i) emerged (ii) failed to develop, and were hyperparasited by (iii) insects or (iv) fungi. Second, we used amplicon sequencing (metabarcoding) to identify to which species belong the 3002 hyperparasitic insects found in all the collected galls. Individuals were pooled by sampling sites and DNA was extracted for each pool using the E.Z.N.A Tissue DNA kit (Omega BIO-TEK) extraction protocol. A Polymerase Chain Reaction (PCR) was performed to amplify the ribosomal Internal Transcribed Spacers 2 (ITS2), commonly used as DNA barcodes and phylogenetic markers in insects. Both forward et reverse primers were obtained from Viviani et al. (99) and were synthetized in 2 different versions with 1N or 3N used as spacers between the Illumina adapter sequence and the marker locus-specific sequence (1N; forward: 5’[TCGTCGGCAGCGTCAGATGTGTATAAGAGACAG]NTGTGAACTGCAGGACACATG 3’, and reverse: 5’[GTCTCGTGGGCTCGGAGATGTGTATAAGAGACAG]NATGCTTAAATTYAG CGGGTA 3’. PCRs were performed in 35 μL reaction volume, containing ∼ 20 ng DNA template, 0.1 μM of each dNTP, 0.04 μM of each primer and 0.7 U Phusion High-Fidelity DNA polymerase (FINNZYMES OY, Espoo, Finland). The thermal profile of the PCR was as follows: initial denaturation at 98°C for 30 seconds, followed by 16 to 22 cycles (depending on the sample) of [denaturation at 98°C for 10 seconds, locus-specific annealing at 52,7°C for 30 seconds, elongation at 72°C for 18 seconds], and a final elongation at 72°C for 10 minutes. The PCR products were checked on 1% agarose gel. Libraries were then generated using Nextera index and Q5 high fidelity DNA polymerase (New England Biolabs). PCR products were normalized with SequalPrep plates (Thermofischer). Paired-end sequencing was performed at the Bio-Environment platform (University of Perpignan Via Domitia Perpignan, France) on a MiSeq system (Illumina) using 2×300bp v3 chemistry. Sequences obtained from the 44 samples (24 samples from 2019 and 20 samples from 2020) have been deposited in the NCBI Sequence Read Archive (SRA) under the BioProject accession number PRJNA1111469 and were processed with the FROGS pipeline (100) available on the Genotoul Bioinfo galaxy server (101). Clustering step was made using SWARM (102) with aggregation distance of d=1. Only OTUs with at least 50 sequences were conserved and manually affiliated using blastn based on the NCBI database best blast hit. Taxonomic identifications were accepted for OTUs sequences with at least 90% of query cover and identity with the blast subject, but for *D. kuriphilus* sequences whose NCBI shorter reference sequences only allow for 86% of query cover (Data S2). A phylogenetic tree of these OTUs and reference sequences of parasitoids previously identified in chestnut tree galls in the South of France (50) (*Torymidae, Eurytomidae, Eupelmidae*) was then built using the Geneious MAFFT alignment and tree builder with default parameters (Fig. S1). Third, to quantify the proportion of *T. sinensis* among all hyperparasitic insects, 45 galls were collected in each of our 8 sampling stations and brought to the laboratory for an emergence experiment. The 360 collected galls were distributed among 72 pots closed with microporous tape to allow gas exchange. Emergence of adult hyperparasites was followed from October 2019 to May 2020 by collecting emerging individuals every week and placing each of them in an individual 0.5mL Eppendorf containing 200μL of alcohol at 100% and stored at -20°C. Morphological identification then allowed to discriminate between *T. sinensis* and other insect species, and therefore to estimate the relative rates of infection by the introduced and native hyperparasitic insects.

### Dynamical modelling of the C. sativa - D. kuriphilus - hyperparasites interactions

We adapted the seminal host-parasitoid model proposed by Nicholson and Bailey (103) to describe the interactions between *C. sativa, D. kuriphilus* and its hyperparasites. The core of our modelling was made of formal representations of the life cycles of *D. kuriphilus* and its hyperparasitic control agent, *T. sinensis*, as described in section ‘*Dryocosmus kuriphilus* life-history and its bottom-up and top-down determinants’, to allow predicting the yearly dynamics of the number of (*T. sinensis*) parasitoid attempting to lay their eggs into (*D. kuriphilus*) host larvae in (the spring of) year t, which we refer to as P(t) and H(t), respectively (Fig. 7). The *D. kuriphilus* host population is first split into larvae escaping parasitism and those who do not, according to a function F_s_ that gives the proportion of non-parasitized larvae with respect to the abundance and searching efficacy of *T. sinensis* in a forest with a proportion pc of chestnut trees (see below, section ‘Modelling ‘top-down’ control’). These two sets of parasitized and non-parasitized individuals initiate the built-up of the next year populations of *T. sinensis* and *D. kuriphilus*.

**Fig. 7.**
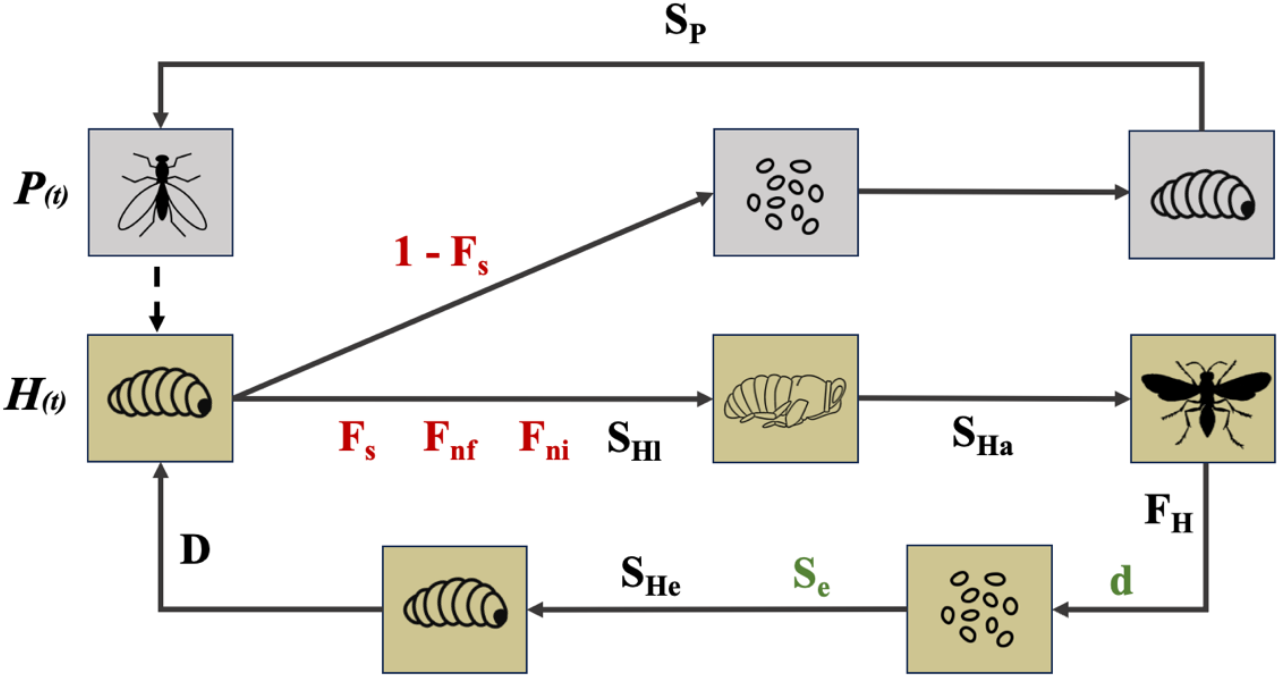
Modelling of the *D. kuriphilus* life-cycle and its bottom-up and top-down control. The lower part of the graph summarizes the different stage of *D. kuriphilus* life-cycle and the effects of the hyperparasitic insects and fungi (top-down control, red) and of the tree species and genetic susceptibility (bottom-up control, green) on *D. kuriphilus* survival and reproduction. The upper part of the graph describe the life-cycle of the introduced control agent, *T. sinensis*. All parameters and functions (D, d, S_e_ and F_s_) are defined in the main text and their estimates provided in Table S2 and Supplementary Text S2.

Larvae non-parasitized by *T. sinensis* face additional challenges by native hyperparasite fungi and insects that were modelled as constant pressures since such populations typically grow on their local host community. *D. kuriphilus* larvae escape those challenges with probabilities denoted F_nf_ and F_ni_, respectively. Larvae escaping all hyperparasites develop at rate S_Hl_ into pupae that ultimately moult into adults with probability S_Ha_. The emerging adults produce on average F_H_ eggs per individual and a fraction d of those eggs are successfully deposited in chestnut tree buds. According to our GLMM analysis of the ‘bottom-up’ control of *D. kuriphilus*’ (Table 1A-B), the fraction d was set to increase with the frequency of chestnut trees (see below, section ‘Modelling ‘bottom-up’ control’). To develop into larvae and enter an overwintering dormant stage, the deposited eggs must survive to the tree hypersensitive response triggered by oviposition and to intrinsic causes of mortality, which they do with probabilities denoted S_e_ and S_He_, respectively. To account for the effect of individual tree genetic differentiation (g) on the level of *D. kuriphilus* infestation (Table 1C), the survival probability S_e_ was defined with respect to the distribution of such genetic differentiation and its effect on the individual rate of infestation (see below, section ‘Modelling ‘bottom-up’ control’). Finally, to complete the description of *D. kuriphilus* life-cycle, we modelled a density dependent survival of dormant larvae, D, as an increase in the number of larvae per bud is known to result in intraspecific competition for space during chambers formation and to the death of first instar larvae (see below, section ‘Modelling the density-dependent regulation of the *D. kuriphilus* larvae population’). The complementary part of *D. kuriphilus* larvae, i.e. the fraction 1-F_s_ not escaping parasitism by *T. sinensis*, is set to die as the hyperparasite larvae feed on them to become the next generation of adults with probability S_P_. All these processes combined into the following set of non-linear difference equations describing the yearly dynamics illustrated in Fig. 7 :

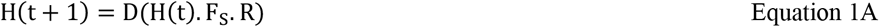

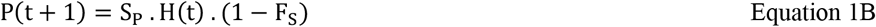

where H(t). F_S_. R corresponds to the number of *D. kuriphilus* dormant larvae, and R = *F*_*ni*_. *F*_*nf*_. *S*_*Hl*_. *S*_*Ha*_. *F*_*H*_. *d*. *Se*.. *S*_*He*._.

The definition of functions D, d, S_e_ and F_S_ were completed by modelling further the biological mechanisms underlying the density-dependent survival of *D. kuriphilus* larvae and the bottom-up and top-down control of its population dynamic. This was done by accounting for both the descriptions of these mechanisms in the literature, and the ecological and genomic analyses intended in this study.

### Modelling the density-dependent regulation of the D. kuriphilus larvae population

A decrease of the within gall survival with larval population density has been observed for *D. kuriphilus* (104) and for other plants-gall-forming parasite interactions (105). Such a decrease comes from a basic form of competition for space where winning larvae get all the resource required to make their chamber, while the other ones die. This typical ‘contest’ competition (106) can be described using a derivation of the Skellam model (107) proposed by Brännström and Sumpter (108):

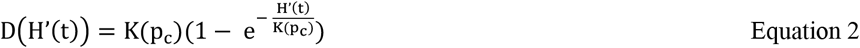

where *H*^′^(t) = *H*(t). F_S_. R denotes the number of *D. kuriphilus* dormant larvae and *K*(p_c_) stands for the maximal amount of larvae that can be sustained, which was set to increase linearly with the proportion p_c_ of chestnut-trees until it reaches a maximal value K estimated from field data (Table S2) for a forest made of only chestnut trees, i.e. *K*(p_c_) = *K*. p_c_.

### Modelling ‘bottom-up’ control

The ‘bottom-up’ determinants of *D. kuriphilus* control naturally occur at two stages of its life-cycle; the oviposition on chestnut buds and the development of eggs into first larvae stage (Fig. 7), which was included in our modelling as described below.

#### The effect of the frequency of chestnut trees on D. kuriphilus’ oviposition rate

Our GLMM analyses have shown that the rates of *D. kuriphilus’* prevalence and infestation are positively correlated with the frequency of chestnut trees in a forest patch. Such an effect had already been documented and associated with an increase of the oviposition rate of *D. kuriphilus* in chestnut tree buds (40). The proportion of eggs deposited on buds was then modelled with respect to the chestnut trees frequency p_c_ and *D. kuriphilus* preference for its resource, *α*_*c*_, using a typical ‘host-choice’ function:

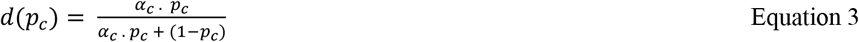

which, according to the low ability of *D. kuriphilus* females to navigate towards chestnut trees in the wild (i.e. *α*_*c*_ = 1, Table S2), ultimately led us to consider d as directly proportional to p_c_.

### The effect of chestnut tree genetic differentiation on D. kuriphilus’s eggs development rate

The susceptibility of chestnut trees to *D. kuriphilus* infection has been shown to vary between individuals in wild populations (36), which we confirmed and further associated with individual genetic differentiation in the chestnut tree populations of the Pyrénées-Orientales (see Table 1). While such variations could hypothetically result from heterogeneities in oviposition or in *D. kuriphilus* development inside buds, it has been shown that females do not display oviposition preferences between resistant and susceptible varieties (109). Instead, an hypersensitive response, a key resistance mechanism against galling insects (110), has been documented on resistant trees with tissue modifications induced by egg secretions soon after oviposition (56) and the accumulation of H_2_O_2_ in the inner part of infested buds (61). Such early hypersensitive reactions in resistant trees are highly consistent with the observed failure of egg development into first instar larvae within chestnut tree buds of resistant varieties (104). Accordingly, the proportion of eggs surviving to the tree hypersensitive response was modelled as the weighted average of the individual tree susceptibility to *D. kuriphilus* infestation defined with respect to its level of genetic differentiation, s(g), where weights were set with respect to the distribution of such individual genetic differentiation, f(g), which led to:

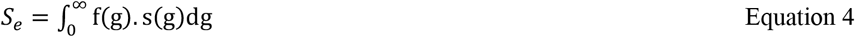

A model selection approach based on the infestation and genetic data collected in this study allowed for the identification of functions *f*(*g*) and *s*(*g*) as 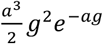 and 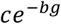, respectively (Supplementary Text S2). The subsequent integration of their product led to *S*_*e*_ = *ca*^3^(*a* + *b*)^−3^. Using this expression and the values of a, b and c inferred while fitting the shape of function s(g) and f(g) to our field data (Table S4 in Supplementary Text S2), we derived the local estimate of the average chestnut tree susceptibility to *D. kuriphilus*, S_e_, for the Pyrénées-Orientales.

### Modelling ‘top-down’ control

The larvae of *D. kuriphilus* typically suffer mortality inflicted by naturally occurring enemies, i.e. hyperparasitic insect species and fungi (42, 67) and by its introduced biological control agent, *T. sinensis* (83), that all contribute to their ‘top-down’ control. The forces of infection exerted by native enemies were modelled as constant according to our field data (Fig. 3 and Table S2), while the force of infection exerted by *T. sinensis* was modelled explicitly according to the dynamical interaction between *D. kuriphilus* and its specific control agent. This force, represented by 1 - F_s_, was derived under the typical assumptions that encounters occur randomly and that once a *D. kuriphilus* larvae has been parasitized, additional encounters will not increase the number of deposited eggs. The proportion of *D. kuriphilus* larvae escaping parasitism was then modelled using a Poisson distribution, which led to:

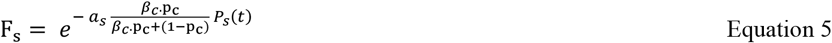

where *a*_*S*_ and *β*_*C*_ represent the search area of *T. sinensis* and its preference for chestnut trees. Since the available data show a low ability of *T. sinensis* adults to fly towards galls of *D. kuriphilus* (i.e., *β*_*C*_ = 1, Table S2), their encounter rate was set to be directly proportional to p_c_.

Lumping together equations 1 to 5, while accounting for the simplifications arising from the low ability of both *D. kuriphilus* and *T. sinensis* to navigate towards their hosts, i.e. *α*_*c*_ = 1 and *β*_*c*_ = 1, we obtained the following modelling framework that we used to integrate our field and genomic data to further assess the dynamics of *D. kuriphilus* invasion and its bottom-up and top-down control:

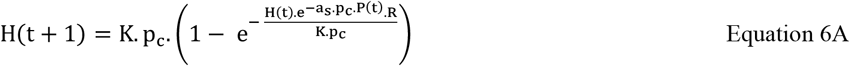

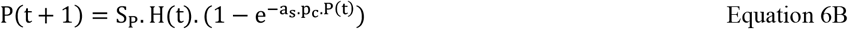

With R = *F*_*ni*_. *F*_*nf*_. *S*_*Hl*_. *S*_*Ha*_. *F*_*H*_. *p*_*c*_. *S*_*e*_ *S*_*He*_

### Model predictions about D. kuriphilus invasion dynamics and its top-down and bottom-up control

We first used our modelling to quantify the invasion potential of *D. kuriphilus* in the Pyrénées-Orientales. We derived the expression of *D. kuriphilus* per capita annual multiplication rate, 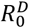, from equation 6A (See Supplementary Text S1), and calculated its value according to the estimates of i) *D. kuriphilus* life-history parameters (Table S2), ii) the average frequency and genetic susceptibility of chestnut trees observed in our field study (i.e. 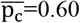 and 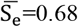), and iii) the average forces of infection exerted by native hyperparasitic insects (i.e.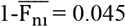) and fungi (i.e.,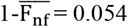). To further quantify the impact of the forest environment and of the native hyperparasite community on *D. kuriphilus* invasion potential, we performed a twofold sensitivity analysis of the estimate of 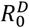 to changes in the values of p_c_, S_e_, 1-F_ni_ and 1-F_nf_. We evaluated i) the linear rate of variation of 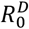 in response to 1% changes in the estimates of each of these parameters and ii) the ranges of variations of 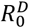 that are expected from the variations observed in these parameters between our 8 sampling stations. We then investigated the dynamics of interaction between *D. kuriphilus* and its control agent, *T. sinensis*. We performed typical local stability analysis of the system of difference equations 6A-B (111) in order to identify the conditions allowing for i) *D. kuriphilus* to be able to spread, ii) *T. sinensis* to spread and persist in a stable equilibrium of coexistence with *D. kuriphilus*, and for iii) the spread of *T. sinensis* to lead to stable oscillations between *D. kuriphilus* and *T. sinensis*. This allowed to assess the expected outcomes of the introduction of the control agent in different forest environments characterized by their frequency and genetic susceptibility of chestnut trees. While the conditions for *D. kuriphilus* and *T. sinensis* to spread could be established analytically, those for the introduction of

*T. sinensis* to lead to either a stable coexistence or oscillations with *D. kuriphilus* were identified numerically by running simulations with R 4.3 (90). The model was then run for 100 generations after the introduction of 1000 *T. sinensis* adult individuals in a forest environment where *D. kuriphilus* has reached its carrying capacity, i.e. with H(0) = K. The independence of the outcome of the interaction with respect to initial conditions was tested by further considering the introduction of *T. sinensis* in an early stage of *D. kuriphilus* invasion, i.e. with H(0) = K/10, and by setting the amount of introduced *T. sinensis* individuals to 100, 1000 and 10000.

Finally, we used similar simulations to assess the potential efficacy of control interventions based on the release of *T. sinensis* according to the frequency and genetic susceptibility of chestnut trees in the forest environment. Simulations were run with and without the control agent, and the effectiveness of the control intervention was measured as the percentage of reduction in the average abundance of *D. kuriphilus*. Such measurement was carried out for the range of p_c_ and S_e_ values allowing for the spread of *T. sinensis*, which we identified as explained above.

## ACKNOWLEDGMENTS

We are grateful to Jean-François Allienne, Margot Doberva and Michèle Laudié from the Bio-Environment platform (UPVD, Région Occitanie, CPER 2007-2013 Technoviv, CPER 2015-2020 Technoviv2) for technical support in library preparation and sequencing.

## Author contributions

Conceptualization: JLZ, SB, RR. Methodology: JLZ, SB, RR, ET. Investigation: JLZ, RR. Visualization: JLZ, RR. Funding acquisition : JLZ, SB. Supervision: SB, ET, JB. Writing—original draft: JLZ, SB. Writing—review & editing: All authors.

## Data and materials availability

All data needed to evaluate the conclusions in the paper are present in the main text and/or the Supplementary Materials. RAD-seq sequences of the 96 genotyped chestnut trees are available in the NCBI Sequence Read Archive (SRA) under the BioProject accession number PRJNA1111809 and metabarcoding sequences of the 3002 hyperparasitic insects collected in chestnut tree galls have been deposited in the NCBI Sequence Read Archive (SRA) under the BioProject accession number PRJNA1111469.

