## Supplementary Materials for "Unravelling the bottom-up and top-down control of a worldwide chestnut tree pest invader through integrative ecological genomics"

### SUPPORTING INFORMATION

**Supplementary Text. Section 1**

Derivation of the per capita annual multiplication rate of the invasive *Dryocosmus kuriphilus* ($R_0^D$).

The equation 6A in the main text gives the abundance of *D. kuriphilus* in year (t+1):

$$H(t+1) = K \,.\, p_c \,.\left(1 - e^{-\frac{H(t).e^{-a_s.p_c.P_S(t)}.R}{K.p_c}}\right)$$

In the absence of the control agent, *T. sinensis*, i.e. when $P_s(t) = 0$, it becomes:

$$H(t+1) = K \,.\, p_c \,.\left(1 - e^{-\frac{H(t).R}{K.p_c}}\right)$$

Further focusing on the early stage of a local invasion, the abundance of *D. kuriphilus* is typically low so that $\frac{H(t).R}{K.p_c}$ tends toward 0. Using $\lim_{x \to 0} 1 - e^{-x} \approx x$, the above equation further simplifies to:

$$H(t+1) \approx K \,.\, p_c \,.\frac{H(t).R}{K.p_c} = H(t).R$$

In the early stage of a *D. kuriphilus* invasion, while *T. sinensis* has not yet been introduced to control its spread, the per capita annual multiplication rate of the invasive is thus equal to:

$$R_0^D = \frac{H(t+1)}{H(t)} = R$$

with $R = F_{ni} \,.F_{nf} \,.S_{Hl} \,.S_{Ha} \,.F_H \,.p_c \,.S_e \,.S_{He}$, as shown while deriving equations 1A and 1B.

**Supplementary Text. Section 2.**

Model selection approach for the identification of functions f(g) and s(g).

*Distribution of chestnut tree individual genetic divergence: f(g).*

We considered three models $f_m(g)$ to be fitted to the observed distribution of individual genetic divergence. These models allow describing three competing hypotheses about the density of probability of chestnut trees carrying a genetic divergence g, that can decrease monotonically according to $f_1(g) = e^{-ag}$ (m=1) or $f_2(g) = e^{-ag^2}$ (m=2), or to increase before to decrease according to $f_3(g) = g^n e^{-ag}$ (m=3), which corresponds to a standard Gamma model when n = 1. The second model was chosen to allow for a larger number of individuals with low divergence than the first, while the third model provides a non-monotonic density probability function with a maximum at an intermediate level of genetic distance. Each of the above functions was normalized to ensure that its integral over the range of possible values of genetic divergence, i.e. from 0 to infinity, is equal to 1. This led to the following expressions:

(m=1): $f_1(g) = \int_0^\infty e^{-ag} = ae^{-ag}$

(m=2): $f_2(g) = \int_0^\infty e^{-ag^2} = 2\sqrt{\frac{a}{\pi}}e^{-ag^2}$

(m=3; n=1): $f_3(g) = \int_0^\infty ge^{-ag} = a^2ge^{-ag}$

(m=3; n=2): $f_3(g) = \int_0^\infty g^2e^{-ag} = \frac{a^3}{2}g^2e^{-ag}$

(m=3; n=3): $f_3(g) = \int_0^\infty g^3e^{-ag} = \frac{a^4}{6}g^3e^{-ag}$

(m=3; n=4): $f_3(g) = \int_0^\infty g^4e^{-ag} = \frac{a^5}{24}g^4e^{-ag}$

The expected probability for a given tree individual to carry a genetic divergence g was then assumed to follow a Poisson distribution with mean parameter: $\lambda = f_m(g)$, which we used in the model selection process described below.

*Relationship between infestation (galls count) and individual genetic divergence: s(g).*

We considered a probabilistic model of infestation assuming that each individual tree receives a random number of attempts to lay a set of eggs (forming a gall) by *D. kuriphilus*, and that the rate of success of each of those attempts depends on the tree genetic divergence g. We then used similar distributions as for the identification of f(g). The second model; $s_2(g) = e^{-bg^2}$, allowed for a larger class of the chestnut tree population (with low levels of genetic divergence) to be highly susceptible to infection than under the first (exponential) model given by $s_1(g) = e^{-bg}$. The non-monotonous model; $s_3(g) = g^n.e^{-bg}$, allowed for individuals with intermediate levels of genetic divergence to be the most susceptible to infection. Under each model (m=1,2,3), the expected number of galls observed on a given tree individual follows a Poisson distribution with mean parameter $\lambda = cs_{\mathrm{m}}(\mathrm{g})$, where c stands for the adjusted per tree mean number of oviposition attempts made by *D. kuriphilus*, which we used in the model fitting and selection approach described below.

*Model fitting and selection.*

Each of the functions $f_1(g)$ to $f_3(g)$ and $s_1(g)$ to $s_3(g)$ were fitted to the distribution of genetic divergence calculated from the 92 individual chestnut tree genomes and to the observed relationship between their rate of infestation by *D. kuriphilus* and level of genetic divergence, respectively. The fits were achieved by maximizing the log-likelihood ($LLH_m$) with respect to the set of model parameters ($\theta_m$).

The maximum likelihood estimator of the parameter a could be identified analytically for functions $f_1(g)$ and $f_2(g)$. For m=1, the log-likelihood function was defined as $\mathrm{LLH}_1(a) = \sum_{i=1}^{n} \log ae^{-ag_i} = n \log a - a \sum_{i=1}^{n} g_i$, where $g_i$ denotes the rate of private alleles of the $i^{th}$ individual in a sample of size n. This log-likelihood function was then differentiated with respect to a ; $\frac{\partial LLH_1(a)}{\partial a} = -n\frac{\partial \ln a}{\partial a} - \sum_{i=1}^{n} g_i \frac{\partial a^{-1}}{\partial a} = -\frac{1}{a^2}(na - \sum_{i=1}^{n} g_i)$, and the estimator of a was found by solving $\frac{\partial LLH_1(a)}{\partial a} = 0$, which led to :

$$\hat{a} = \frac{1}{\bar{g}}$$

where $\bar{g}$ stands for the average rate of individual genetic divergence.

For m=2, the log-likelihood function was defined as $\mathrm{LLH}_2(a) = \sum_{i=1}^{n} \log 2\sqrt{\frac{a}{\pi}} e^{-ag_i^2} = n \log 2\sqrt{\frac{a}{\pi}} - a\sum_{i=1}^{n} g_i^2$ and differentiated with respect to a ; $\frac{\partial \mathrm{LLH}_2(a)}{\partial a} = n\frac{\partial \ln 2\sqrt{\frac{a}{\pi}}}{\partial a} - \sum_{i=1}^{n} g_i^2 = \frac{n}{2a} - \sum_{i=1}^{n} g_i^2$, which led to :

$$\hat{a} = \frac{1}{2\overline{g^2}}$$

where $\overline{g^2}$ stands for the average rate of the square of individual genetic divergence.

The maximum likelihood estimates of the parameter a of model $f_3(g)$ and $s_1(g)$ to $s_3(g)$were identified numerically using R 4.1.1 (90). For both the observed distribution of genetic divergences and the observed relationship between infestation and genetic divergences, the best (fitted) model was then selected using the standard Akaike Information Criterion, i.e. $\mathrm{AIC}_m$= 2p - 2$\mathrm{LLH}_m$, where p stands for the number of model parameters, and we further calculated the associated weights of Akaike ($w_m$) for each model.

The outcomes of the model fitting and selection are summarized in table below.

| **Model** | **Maximum log-likelihood ($\mathrm{LLH}_m$)** | **Model parameters estimates** | **Akaike Information Criterion ($\mathrm{AIC}_m$)** | **Weight of Akaike ($w_m$)** |
|---|---|---|---|---|
| *Distribution of chestnut tree individual genetic divergence: f(g).* | | | | |
| m=1 | 818.1238 | $a$ = 19784.41 | -1634.248 | 2.103e-10 |
| m=2 | 820.7526 | $a$ = 195711415 | -1639.505 | 2.9136e-09 |
| m=3 (n=1) | 837.5209 | $a$ = 39568.82 | -1671.042 | 0.0205 |
| m=3 (n=2) | 841.2873 | $a$ = 59353.2 | -1678.575 | 0.8878 |
| m=3 (n=3) | 839.0124 | $a$ = 79137.6 | -1674.025 | 0.0913 |
| m=3 (n=4) | 833.5165 | $a$ = 98922.2 | -1663.033 | 0.0004 |
| *Relationship between the rate of infestation and individual genetic divergence: s(g).* | | | | |
| m=1 | -1175.094 | $b$ = 9815.96<br>$c$ = 1.0769 | 2354.188 | >0.9999 |

| | | | | |
|---|---|---|---|---|
| m=2 | -1192.467 | $b = 0.7982$<br>$c = 48317500$ | 2388.934 | 2.851e-08 |
| m=3 (n=1) | -1191.903 | $b = 63930$<br>$c = 28810$ | 2389.806 | 1.8435e-08 |

**Table S4.** Summary table of each model parameters estimates, their maximum log-likelihood, Akaike information criterion and weights of Akaike for the f(g) and s(g) function.

The weights of Akaike calculated after the fit of the competing functions f(g) to the observed distribution of genetic divergence showed a strong support for the third model with n=2, i.e. $f_3(g) = \frac{a^3}{2} g^2 e^{-ag}$, as its weight indicated that there is 88.8% chance that it is the best approximating model. The fit of the functions s(g) to the observed relationship between infestation and genetic led to an even stronger support for the first model, i.e. $s_1(g) = ce^{-bg}$, with a <99% chance that it represents best the data.

The selection of functions $f(g) = \frac{a^3}{2} g^2 e^{-ag}$ and $s(g) = ce^{-bg}$ subsequently allowed to get an estimator of the proportion of eggs surviving to the tree hypersensitive response ($S_e$). As explained in the main text, $S_e$ can be defined as the average of the individual tree susceptibility to *D. kuriphilus* infestation (defined with respect to its level of genetic differentiation); s(g), weighted by the distribution of such individual genetic differentiation, f(g). Using the selected functions $f(g)$ and $s(g)$, the expression of $S_e$ (equation 4) becomes:

$$S_e = \frac{a^3 c}{2} \int_0^{\infty} g^2 e^{-(a+b)g} dg$$

Integrating by parts twice allows to obtain the estimator of $S_e$:

$$\widehat{S_e} = a^3 c (a+b)^{-3}$$

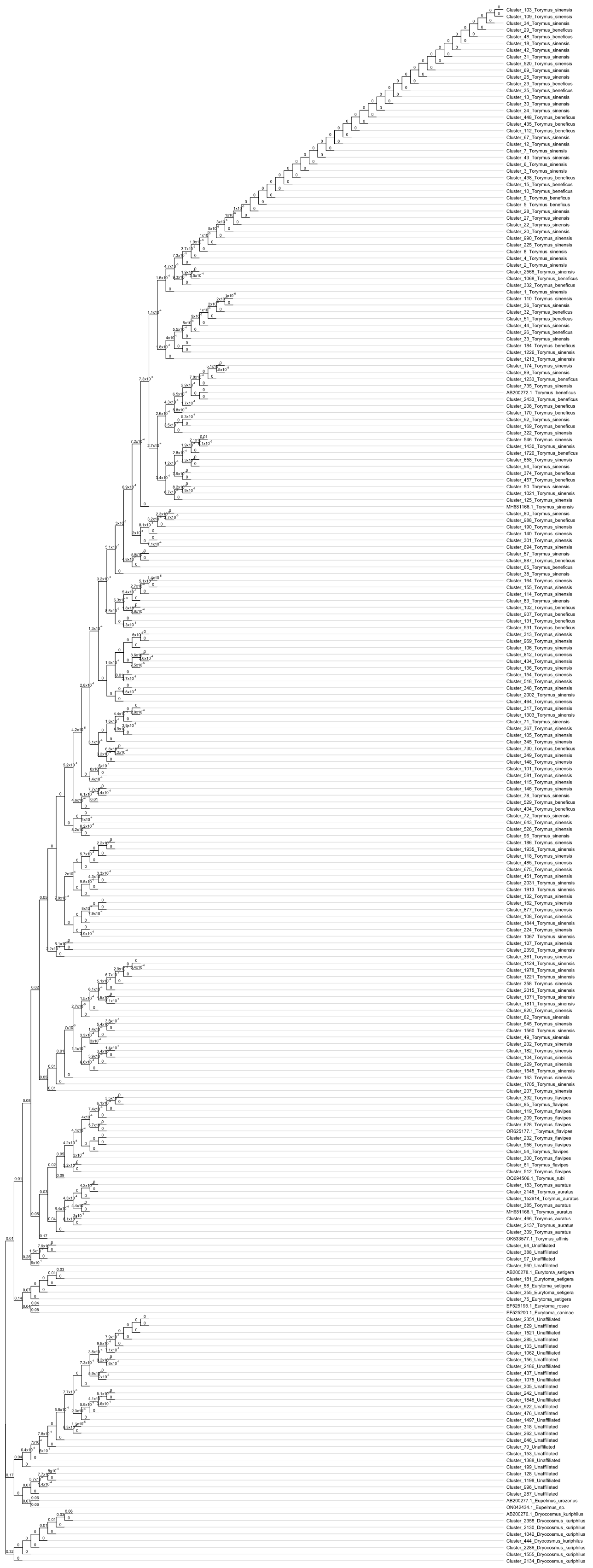

**Fig. S1.** Phylogenetic tree of sequences corresponding to the 219 OTUs identified from the metabarcoding analysis.

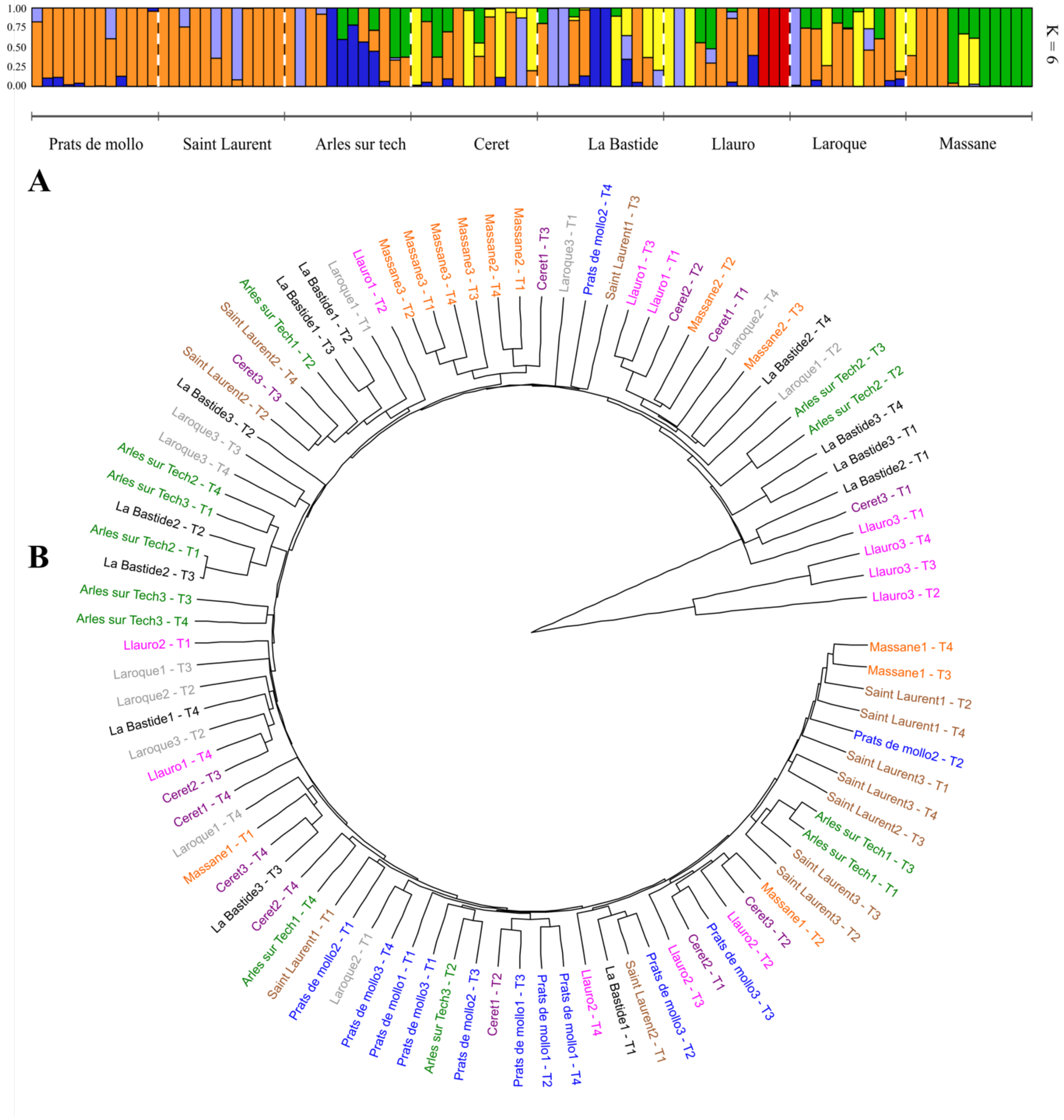


**Fig. S2.** Genetic structure of the chestnut tree populations in the Pyrénées-Orientales identified from 95 individuals sequenced across our 8 sampling stations (see section 'Material and Method - *Genetic structure of the C. sativa populations*'). Colors in figure A show the 6 putative ancestral populations identified, while colors in figure B are set to differentiate our 8 sampling stations (according to the color code used in Figure 2A). This initial analysis showed that individuals from the third sampling site located in Llauro (Llauro3 - T2-3-4 in figure B) were genetically identical and very divergent from the rest of the sequenced individuals, so that they were treated as outliers and this sampling site was removed from analyses leading to figures 1-2 and table 1.

| Stations | Chestnut tree resource | | | *D. kuriphilus* | | | | Native parasitoïd impact | | |
|---|---|---|---|---|---|---|---|---|---|---|
| | Density | Frequency (pc) | Genetic susceptibility (Se) | Prevalence rate | | Infestation rate | | Hymenopteran (1-Fni) | Fungi (1-Fnf) | |
| | | | | 2019 | 2020 | 2019 | 2020 | 2019 | 2019 | 2020 |
| Vallespir | 1055 | 0.639 | 0.635 | 0.845 | 0.472 | 0.069 | 0.023 | 0.036 | 0.065 | 0.053 |
| Prats de Mollo | 1077 | 0.731 | 0.707 | 0.867 | 0.173 | 0.022 | 0.002 | 0.068 | 0.049 | 0.063 |
| Site 1 | 1120 | 0.491 | 0.720 | 0.780 | 0.040 | 0.014 | 0 | 0.273 | 0.036 | 0.133 |
| Site 2 | 870 | 0.908 | 0.639 | 0.940 | 0.260 | 0.029 | 0 | 0 | 0.039 | 0.051 |
| Site 3 | 1240 | 0.823 | 0.768 | 0.880 | 0.220 | 0.0236 | 0.005 | 0 | 0.070 | 0.061 |
| Saint Laurent | 1277 | 0.671 | 0.671 | 0.886 | 0.473 | 0.112 | 0.022 | 0 | 0.086 | 0.074 |
| Site 1 | 1030 | 0.418 | 0.535 | 0.714 | 0.020 | 0.066 | 0 | 0 | 0.106 | 0 |
| Site 2 | 1230 | 0.854 | 0.731 | 0.980 | 0.560 | 0.144 | 0.035 | 0 | 0.047 | 0.093 |
| Site 3 | 1570 | 0.694 | 0.774 | 0.960 | 0.840 | 0.127 | 0.033 | 0 | 0.103 | 0.050 |
| Arles sur Tech | 1173 | 0.580 | 0.526 | 0.818 | 0.513 | 0.062 | 0.023 | 0.049 | 0.081 | 0.031 |
| Site 1 | 1070 | 0.869 | 0.427 | 0.920 | 0.500 | 0.052 | 0.008 | 0 | 0.073 | 0.021 |
| Site 2 | 1330 | 0.286 | 0.587 | 0.703 | 0.600 | 0.057 | 0.013 | 0.111 | 0.108 | 0.025 |
| Site 3 | 1120 | 0.652 | 0.581 | 0.800 | 0.440 | 0.079 | 0.053 | 0 | 0.064 | 0.046 |
| Céret | 693 | 0.591 | 0.650 | 0.807 | 0.727 | 0.081 | 0.04 | 0.030 | 0.046 | 0.054 |
| Site 1 | 740 | 0.581 | 0.686 | 0.900 | 1 | 0.132 | 0.055 | 0.100 | 0.047 | 0.088 |
| Site 2 | 760 | 0.908 | 0.755 | 0.980 | 0.960 | 0.081 | 0.058 | 0 | 0.044 | 0.038 |
| Site 3 | 580 | 0.190 | 0.530 | 0.540 | 0.220 | 0.018 | 0.003 | 0.035 | 0.047 | 0.033 |
| Aspres | 1100 | 0.594 | 0.569 | 0.702 | 0.256 | 0.030 | 0.004 | 0.017 | 0.043 | 0.036 |
| La Bastide | 1157 | 0.504 | 0.612 | 0.63 | 0.033 | 0.025 | 0.003 | 0.020 | 0.026 | 0.060 |
| Site 1 | 1310 | 0.473 | 0.756 | 0.900 | 0.020 | 0.006 | 0.001 | 0 | 0.022 | X |
| Site 2 | 850 | 0.424 | 0.667 | 1 | 0.080 | 0.063 | 0.008 | 0.046 | 0.018 | 0.060 |
| Site 3 | 1310 | 0.588 | 0.455 | 0.100 | 0 | 0.007 | 0 | 0 | 0.037 | X |
| Llauro | 1015 | 0.739 | 0.511 | 0.800 | 0.590 | 0.037 | 0.005 | 0.015 | 0.056 | 0.034 |
| Site 1 | 1270 | 0.724 | 0.566 | 0.960 | 0.740 | 0.046 | 0.005 | 0 | 0.061 | 0.027 |
| Site 2 | 760 | 0.763 | 0.460 | 0.640 | 0.440 | 0.033 | 0.005 | 0.040 | 0.066 | 0.025 |
| Site 3 | 190 | 0.895 | X | 0.780 | 0.640 | 0.010 | 0.011 | 0 | 0.037 | 0.053 |
| Albères | 753 | 0.400 | 0.593 | 0.565 | 0.130 | 0.031 | 0.002 | 0.096 | 0.096 | 0.035 |
| Laroque | 1073 | 0.562 | 0.575 | 0.634 | 0.147 | 0.035 | 0.003 | 0.152 | 0.054 | 0.037 |
| Site 1 | 1170 | 0.735 | 0.581 | 0.760 | 0.320 | 0.037 | 0.009 | 0.133 | 0.044 | 0.041 |
| Site 2 | 1150 | 0.583 | 0.702 | 0.744 | 0.120 | 0.041 | 0.001 | 0.167 | 0.050 | 0 |
| Site 3 | 900 | 0.311 | 0.466 | 0.366 | 0 | 0.027 | 0 | 0.160 | 0.079 | X |
| La Massane | 433 | 0.300 | 0.611 | 0.492 | 0.110 | 0.027 | 0.001 | 0.042 | 0.144 | 0.030 |
| Site 1 | 250 | 0.200 | 0.687 | 0.393 | 0.074 | 0.011 | 0 | 0.091 | 0.094 | 0 |
| Site 2 | 670 | 0.418 | 0.644 | 0.500 | 0.160 | 0.044 | 0.002 | 0.046 | 0.120 | 0.083 |
| Site 3 | 380 | 0.158 | 0.515 | 0.540 | 0.080 | 0.028 | 0 | 0 | 0.177 | 0.016 |
| Total | 986 | 0.602 | 0.680 | 0.746 | 0.340 | 0.050 | 0.013 | 0.045 | 0.067 | 0.046 |

**Table S1.** Summary table of the invasion of *Dryocosmus kuriphilus* and of its bottom-up and top-down factors assessed in all sampled sites and stations across the three massifs of the Pyrénées-Orientales. The measures include the chestnut tree density, frequency and genetic susceptibility, the prevalence and rate of infestation by *D. kuriphilus*, and the impact of (hymenopteran and fungal) hyperparasites. As described in section 'Material and Methods - *Genetic structure of the C. sativa populations*', three chesnut trees of our third sampling site located in Llauro were almost genetically identical and very divergent from all other sequenced individuals. Accordingly, the measures made in this site (appearing in light grey) were not accounted for in the average values characterizing the chestnut tree resource and its infestation by *D. kuriphilus* in Llauro and in the Vallespir massif.

| | Symbol | Estimate | References and notes |
|---|---|---|---|
| ***Dryocosmus kuriphilus*** | | | |
| Egg intrinsic survival | $S_{He}$ | 0.535 ± 0.003 | (56) [(1)] |
| Egg survival to hypersensitive response | $S_e$ | 0.680 | This study [(2)] |
| Larvae survival | $S_{Hl}$ | 0.936 ± 0.010 | (112) [(3)] |
| Adult survival | $S_{Ha}$ | 0.952 ± 0.010 | (41) [(3)] |
| Fertility | $F_H$ | 90 ± 6 | This study [(4)] |
| Preference for chestnut trees | $\alpha_c$ | 1 | (113) [(5)] |
| Carrying capacity per hectare | K | 14 880 000 | This study [(6)] |
| ***Torymus sinensis*** | | | |
| Larvae and adult survival | $S_P$ | 0.913 ± 0.004 | (114) |
| Preference for chestnut trees | $\beta_c$ | 1 | (115) [(7)] |
| Searching area | $a_s$ | $4.4 \times 10^{-7} \pm 0.4 \times 10^{-7}$ | This study [(8)] |
| ***Native hymenoptera and fungi*** | | | |
| Probability to escape hymenopteran infection | $F_{ni}$ | 0.955 ± 0.02 | This study [(9)] |
| Probability to escape fungal infection | $F_{nf}$ | 0.946 ± 0.001 | This study [(9)] |

**Table S2.** Definition and estimates of the parameters of the dynamical model of interaction between *Castanea sativa*, *Dryocosmus kuriphilus* and its control agent *Torymus sinensis* and native hyperparasites.

(1) The intrinsic rate of survival of *D. kuriphilus* eggs (accounting for all causes of mortality independent of the host hypersensitive response) was calculated by averaging two separate estimates derived by Nugnes et al. (56) on resistant and susceptible chestnut trees as no statistical difference was found between them. (2) See Supplementary material 6 for details. Our estimate of the egg survival to the hypersensitive response is similar to the difference of egg development rates observed on sensible and resistant trees by Nugnes et al. (56). While similar numbers of oviposition scars were found on buds of sensible and resistant trees, the average number of developping eggs were 23.8 and 16.6, respectively, corresponding to a differential survival of 16.6/23.8=0.739, close to the estimate of $S_e$ derived from the very different approach intended in SM9. (3) The rate of *D. kuriphilus* survival after hatching, i.e. the product of the larval ($S_{Hl}$) and adult ($S_{Ha}$) survival, is equal to 0.871, and very similar to previous estimates of 0.857 (46) and 0.863 (116). (4) The fertility of *D. kuriphilus* was estimated as $F_H = \frac{Ne*S}{1-S}$, where Ne stands for the daily number of eggs that could potentially be laid, which we estimated to be equal to 30 eggs by dividing the number of eggs that are carried by an emerging individual at the onset of its adult lifetime (117) by the 4 days duration of this adult stage (118), and where S denotes the corresponding daily survival estimated to be equal to 0.75 from a geometric model. (5) The preference of *D. kuriphilus* for chestnut trees, $\alpha_c$, was set to 1, according to a Y-tube olfactometer bioassay performed by Germinara et al. (113), where *Castanea sativa* twigs were found not attractive while displayed in a 30cm range from the insect. (6) The *D. kuriphilus* larvae carrying capacity, K, was calculated as the total number of lodges that can be found in chestnut galls per hectare by multiplying the maximum density of chestnut buds per hectare of a pure chestnut tree stand (i.e. when $p_c$=1), estimated to be $2*10^6$ $buds.h^{-1}$ (119), by the maximum number of galls per bud, observed to reach 1.24 $gall.bud^{-1}$ (36) and by the maximum number of lodges that we recorded by gall, i.e. 6. (7) The ability of *T. sinensis* to locate *D. kuriphilus* galls on chestnut trees in a mixed forest environment is expected to be limited as Graziosi and Rieske (115) showed that both olfactory and visual clues are necessary for this hyperparasite to detect its resource in a typically short-range corresponding to the performed Y-tube

olfactometer experiment. (8) The *T. sinensis* searching area, $a_s$, was estimated by deriving its expression from equation 6B describing *T. sinensis* dynamics while considering a typical geometric growth soon after its release, i.e. $P(t+1) = R_0^T(t)P(t)$ where $R_0^T$ denotes *T. sinensis* intrinsic annual growth rate. Straightforward algebraic manipulation lead to $a_s = \frac{1}{p_c.P(t)} \log\left(1 - \frac{R_0^T.P(t)}{S_P.H(t)}\right) P(t)$. The estimate of $a_s$ was then calculated using a value of $R_0^T$ that we derived from Muru et al. (120) by fitting the annual geometric growth of *T. sinensis* parasitism of cynips galls observed over 4 to 6 years in 9 separate orchards, and by retaining the geometric mean of the 32 resulting values as well as their standard deviation. (9) The forces of infection by native species of hyperparasitic insects and fungi were estimated to reach 4.46% and 5.44%, respectively, after dissection of 2110 galls of *D. kuriphilus* and found very consistent across the different localities of our field study, which allowed to calculate the fraction of larvae escaping each of those two taxa of native hyperparasites, i.e. $F_{ni}$ and $F_{nf}$, as their complementary percentages.

| Sample name | Raw reads | Adapter clipped reads | Restriction enzyme filtered reads | All sites | Variant sites | Polymorphic sites | Private alleles | Mean coverage |
|---|---|---|---|---|---|---|---|---|
| Llauro1 - T1 | 4 871 110 | 4 871 008 | 4 870 962 | 8068748 | 104420 | 19762 | 515 | 45,28 |
| Llauro1 - T2 | 5 045 302 | 5 045 224 | 5 045 169 | 8175447 | 107889 | 22554 | 725 | 46,28 |
| Llauro1 - T3 | 5 215 514 | 5 215 367 | 5 215 310 | 8615152 | 102064 | 18375 | 315 | 45,40 |
| Llauro1 - T4 | 4 753 022 | 4 752 943 | 4 752 877 | 7957245 | 98609 | 17175 | 250 | 44,80 |
| Llauro2 - T1 | 1 026 280 | 1 026 268 | 1 026 240 | 4623282 | 59472 | 7897 | 457 | 16,65 |
| Llauro2 - T2 | 1 579 004 | 1 578 982 | 1 578 966 | 4662968 | 69106 | 10625 | 275 | 25,40 |
| Llauro2 - T3 | 6 351 982 | 6 351 859 | 6 351 792 | 13107610 | 134088 | 26945 | 1300 | 36,35 |
| Llauro2 - T4 | 3 705 489 | 3 705 412 | 3 705 377 | 6348059 | 94695 | 17828 | 359 | 43,78 |
| Llauro3 - T1 | 512 414 | 512 407 | 512 402 | 3168063 | 47101 | 6141 | 230 | 12,13 |
| Llauro3 - T2 | 1 192 238 | 1 192 231 | 1 192 220 | 7690913 | 85701 | 28966 | 13245 | 11,63 |
| Llauro3 - T3 | 144 034 | 144 033 | 144 032 | 1740072 | 21405 | 3240 | 503 | 6,21 |
| Llauro3 - T4 | 339 561 | 339 560 | 339 552 | 2572928 | 33658 | 4756 | 2541 | 9,90 |
| Ceret1 - T1 | 5 495 841 | 5 495 735 | 5 495 647 | 10214965 | 111660 | 20274 | 594 | 40,35 |
| Ceret1 - T2 | 5 288 349 | 5 288 258 | 5 288 182 | 8012853 | 102881 | 18655 | 236 | 49,50 |
| Ceret1 - T3 | 4 895 809 | 4 895 724 | 4 895 655 | 7930763 | 107036 | 18099 | 297 | 46,30 |
| Ceret1 - T4 | 1 398 113 | 1 398 079 | 1 398 063 | 5519330 | 75750 | 14327 | 128 | 19,00 |
| Ceret2 - T1 | 4 878 781 | 4 878 758 | 4 878 709 | 7606025 | 106184 | 19874 | 160 | 48,11 |
| Ceret2 - T2 | 5 371 926 | 5 371 766 | 5 371 676 | 8219934 | 103584 | 18556 | 240 | 49,01 |
| Ceret2 - T3 | 4 794 279 | 4 794 150 | 4 794 091 | 7923946 | 100698 | 17418 | 247 | 45,38 |
| Ceret2 - T4 | 5 934 731 | 5 934 687 | 5 934 629 | 8732235 | 108955 | 21103 | 265 | 50,97 |
| Ceret3 - T1 | 5 117 345 | 5 117 317 | 5 117 268 | 10359789 | 120667 | 27540 | 1584 | 37,05 |
| Ceret3 - T2 | 4 892 190 | 4 892 029 | 4 891 982 | 8996196 | 111792 | 22821 | 281 | 40,79 |
| Ceret3 - T3 | 4 680 982 | 4 680 910 | 4 680 503 | 8008649 | 107915 | 20915 | 270 | 43,84 |
| Ceret3 - T4 | 5 580 464 | 5 580 356 | 5 580 299 | 8701660 | 109355 | 19175 | 264 | 48,10 |
| Massane1 - T1 | 5 737 190 | 5 737 114 | 5 737 067 | 8624809 | 108551 | 21332 | 290 | 49,89 |
| Massane1 - T2 | 4 761 474 | 4 761 324 | 4 761 273 | 7971355 | 105249 | 21068 | 208 | 44,80 |
| Massane1 - T3 | 9 081 675 | 9 081 618 | 9 080 884 | 11518764 | 132943 | 28792 | 704 | 59,13 |
| Massane1 - T4 | 5 900 139 | 5 900 003 | 5 899 940 | 8981241 | 115378 | 21964 | 282 | 49,27 |
| Massane2 - T1 | 2 419 985 | 2 419 959 | 2 419 921 | 5258715 | 76633 | 11207 | 323 | 34,51 |
| Massane2 - T2 | 5 855 583 | 5 855 409 | 5 854 851 | 8352030 | 107813 | 20633 | 324 | 52,58 |
| Massane2 - T3 | 5 695 503 | 5 695 353 | 5 695 299 | 8748753 | 108341 | 24907 | 343 | 48,83 |
| Massane2 - T4 | 4 199 211 | 4 199 087 | 4 199 050 | 7564503 | 101457 | 18791 | 282 | 41,63 |
| Massane3 - T1 | 3 899 413 | 3 899 372 | 3 899 348 | 6489558 | 92596 | 14552 | 417 | 45,07 |
| Massane3 - T2 | 3 923 077 | 3 923 023 | 3 922 984 | 6587208 | 95697 | 16205 | 561 | 44,67 |
| Massane3 - T3 | 679 193 | 679 178 | 679 169 | 3518188 | 51440 | 6641 | 121 | 14,48 |
| Massane3 - T4 | 4 437 067 | 4 437 001 | 4 436 941 | 6231498 | 87619 | 14088 | 516 | 53,40 |
| Laroque1 - T1 | 1 402 305 | 1 402 294 | 1 402 278 | 7049395 | 86574 | 15665 | 279 | 14,92 |
| Laroque1 - T2 | 537 673 | 537 673 | 537 629 | 3254139 | 50743 | 7221 | 195 | 12,39 |
| Laroque1 - T3 | 2 684 862 | 2 514 056 | 2 513 835 | 5760271 | 86559 | 12992 | 572 | 34,96 |
| Laroque1 - T4 | 1 363 997 | 1 363 949 | 1 363 937 | 5724743 | 80530 | 13897 | 127 | 17,87 |
| Laroque2 - T1 | 140 433 | 140 433 | 140 432 | 1517145 | 21837 | 2086 | 48 | 6,94 |
| Laroque2 - T2 | 764 665 | 716 410 | 716 400 | 3871969 | 59133 | 8761 | 206 | 14,81 |
| Laroque2 - T3 | 7 948 | 7 432 | 7 432 | 295056 | 3071 | 152 | 8 | 2,02 |
| Laroque2 - T4 | 2 323 896 | 2 178 139 | 2 178 117 | 6862710 | 93141 | 16550 | 217 | 25,40 |
| Laroque3 - T1 | 1 806 892 | 1 806 834 | 1 806 811 | 6879882 | 87188 | 18515 | 205 | 19,70 |
| Laroque3 - T2 | 736 940 | 736 938 | 736 908 | 3216670 | 44609 | 5359 | 340 | 17,18 |
| Laroque3 - T3 | 3 695 156 | 3 695 090 | 3 695 051 | 13747565 | 120243 | 21607 | 1636 | 20,16 |
| Laroque3 - T4 | 2 049 217 | 2 049 182 | 2 049 022 | 8791695 | 102959 | 19236 | 470 | 17,48 |
| La Bastide1 - T1 | 2 074 450 | 2 074 409 | 2 074 391 | 7745863 | 95937 | 19933 | 283 | 20,09 |
| La Bastide1 - T2 | 1 137 146 | 1 069 136 | 1 069 125 | 5181283 | 72727 | 12419 | 94 | 16,46 |
| La Bastide1 - T3 | 2 854 654 | 2 854 562 | 2 854 531 | 8066041 | 102044 | 17100 | 247 | 26,54 |
| La Bastide1 - T4 | 1 915 765 | 1 915 736 | 1 915 704 | 6406598 | 85137 | 13536 | 161 | 22,43 |
| La Bastide2 - T1 | 1 372 490 | 1 372 443 | 1 372 431 | 5371988 | 76565 | 11398 | 146 | 19,16 |
| La Bastide2 - T2 | 103 325 | 103 319 | 103 316 | 1290294 | 18195 | 1984 | 51 | 6,01 |
| La Bastide2 - T3 | 2 142 922 | 2 142 889 | 2 142 872 | 6807685 | 91886 | 17128 | 202 | 23,61 |
| La Bastide2 - T4 | 1 275 268 | 1 275 254 | 1 275 239 | 8242827 | 91904 | 17082 | 501 | 11,60 |
| La Bastide3 - T1 | 3 585 788 | 3 585 781 | 3 585 726 | 5963860 | 87088 | 16086 | 651 | 45,09 |
| La Bastide3 - T2 | 2 318 972 | 2 318 969 | 2 318 937 | 5286875 | 77144 | 12473 | 620 | 32,90 |
| La Bastide3 - T3 | 2 365 738 | 2 365 697 | 2 365 674 | 6649917 | 90862 | 17659 | 114 | 26,68 |
| La Bastide3 - T4 | 1 811 350 | 1 811 294 | 1 811 284 | 6233191 | 83997 | 14980 | 408 | 21,79 |
| Arles sur Tech1 - T1 | 694 135 | 694 133 | 694 127 | 3587204 | 53501 | 9080 | 339 | 14,51 |
| Arles sur Tech1 - T2 | 1 578 365 | 1 578 336 | 1 578 191 | 6465074 | 86905 | 16479 | 275 | 18,31 |
| Arles sur Tech1 - T3 | 1 418 268 | 1 418 249 | 1 418 230 | 5614673 | 78027 | 14847 | 101 | 18,95 |
| Arles sur Tech1 - T4 | 6 083 485 | 6 083 374 | 6 083 315 | 9012920 | 113211 | 23320 | 397 | 50,62 |
| Arles sur Tech2 - T1 | 1 662 905 | 1 662 892 | 1 662 878 | 5807158 | 82352 | 15345 | 144 | 21,48 |
| Arles sur Tech2 - T2 | 4 463 618 | 4 463 535 | 4 463 477 | 8064730 | 103245 | 19068 | 331 | 41,51 |
| Arles sur Tech2 - T3 | 2 171 839 | 2 171 797 | 2 171 621 | 6667109 | 89317 | 16221 | 307 | 24,43 |
| Arles sur Tech2 - T4 | 2 352 143 | 2 352 127 | 2 352 096 | 5586165 | 81055 | 12972 | 514 | 31,58 |
| Arles sur Tech3 - T1 | 1 768 372 | 1 768 332 | 1 768 307 | 5417839 | 70697 | 12736 | 122 | 24,48 |
| Arles sur Tech3 - T2 | 5 912 768 | 5 912 729 | 5 912 235 | 10681745 | 126973 | 25217 | 629 | 41,52 |
| Arles sur Tech3 - T3 | 710 712 | 710 705 | 710 674 | 3244714 | 42783 | 5131 | 316 | 16,43 |
| Arles sur Tech3 - T4 | 5 843 200 | 5 451 070 | 5 451 011 | 8770163 | 104953 | 19861 | 324 | 49,97 |
| Saint Laurent1 - T1 | 8 096 802 | 8 096 633 | 8 096 549 | 11688932 | 131045 | 26896 | 806 | 51,95 |
| Saint Laurent1 - T2 | 5 029 209 | 5 029 111 | 5 029 040 | 14398105 | 123935 | 25298 | 1745 | 26,20 |
| Saint Laurent1 - T3 | 2 808 668 | 2 808 615 | 2 808 590 | 7650588 | 90992 | 16526 | 306 | 27,53 |
| Saint Laurent1 - T4 | 1 669 330 | 1 669 312 | 1 669 292 | 6087694 | 82067 | 15262 | 117 | 20,57 |
| Saint Laurent2 - T1 | 468 037 | 468 030 | 468 027 | 3304789 | 48990 | 6703 | 120 | 10,62 |
| Saint Laurent2 - T2 | 225 553 | 225 553 | 225 548 | 2080162 | 30812 | 3539 | 94 | 8,13 |
| Saint Laurent2 - T3 | 1 800 437 | 1 800 416 | 1 800 392 | 5889332 | 79836 | 15773 | 106 | 22,93 |
| Saint Laurent2 - T4 | 1 574 539 | 1 574 512 | 1 574 498 | 5798839 | 80370 | 15199 | 149 | 20,36 |
| Saint Laurent3 - T1 | 6 918 935 | 6 918 695 | 6 918 624 | 9818379 | 119351 | 24762 | 381 | 52,85 |
| Saint Laurent3 - T2 | 6 410 031 | 6 409 913 | 6 409 845 | 8300609 | 109338 | 18660 | 225 | 57,92 |
| Saint Laurent3 - T3 | 2 407 853 | 2 407 808 | 2 407 794 | 6511095 | 90090 | 16404 | 134 | 27,74 |
| Saint Laurent3 - T4 | 2 345 022 | 2 344 977 | 2 344 938 | 6452623 | 86833 | 16862 | 123 | 27,26 |
| Prats de mollo1 - T1 | 2 515 357 | 2 515 309 | 2 515 290 | 7031553 | 92655 | 19280 | 177 | 26,83 |
| Prats de mollo1 - T2 | 2 322 651 | 2 322 597 | 2 322 390 | 6465477 | 89845 | 15826 | 174 | 26,94 |
| Prats de mollo1 - T3 | 2 222 289 | 2 222 225 | 2 222 193 | 7374923 | 91390 | 16961 | 196 | 22,60 |
| Prats de mollo1 - T4 | 1 654 358 | 1 654 320 | 1 654 297 | 9314180 | 97711 | 18785 | 499 | 13,32 |
| Prats de mollo2 - T1 | 3 767 915 | 3 767 814 | 3 767 788 | 6277521 | 92835 | 14234 | 414 | 45,02 |
| Prats de mollo2 - T2 | 3 900 454 | 3 900 359 | 3 900 319 | 10077251 | 114201 | 19230 | 396 | 29,03 |
| Prats de mollo2 - T3 | 697 450 | 697 448 | 697 434 | 3612363 | 53565 | 7717 | 140 | 14,48 |
| Prats de mollo2 - T4 | 2 245 843 | 2 245 833 | 2 245 815 | 6582657 | 88725 | 14074 | 273 | 25,59 |
| Prats de mollo3 - T1 | 2 637 480 | 2 637 420 | 2 637 398 | 6941260 | 92558 | 15929 | 160 | 28,50 |
| Prats de mollo3 - T2 | 1 865 722 | 1 865 688 | 1 865 517 | 6159856 | 88837 | 15623 | 159 | 22,72 |
| Prats de mollo3 - T3 | 1 675 099 | 1 675 058 | 1 675 047 | 5777968 | 80161 | 16203 | 103 | 21,74 |
| Prats de mollo3 - T4 | 932 604 | 932 590 | 932 585 | 4568062 | 68733 | 11531 | 191 | 15,31 |

**Table S3.** Raw sequencing results for the 96 chestnut trees.

**Data S1. (GWAS-Results.xlsx).** List of loci containing SNPs that were shown to be significantly associated with the rate of chestnut tree infestation by *Dryocosmus kuriphilus* through a GWAS. The GWAS was conducted separately on the infestation phenotypes observed in 2019 and 2020. For each loci, the first and last columns indicate the p-value quantifying the level of association and the year when the SNP p-value was found significant. When an association was found in both years, the lowest of the two resulting p-values is indicated the first column and the corresponding year appears in bold in the last column. The contig where the loci was found, its position, homolog and associated function are reported in the other columns.

**Data S2. (Blast-results-metabarcoding.xlsx).** Summary table of BLAST results for the 219 OTUs identified from the metabarcoding analysis.
